# Systematic Review of Artificial Intelligence use in behavioral analysis of invertebrate and larval model organisms: Methods, Applications and Future Recommendations

**DOI:** 10.1101/2025.10.16.682789

**Authors:** Zuzanna Stępnicka, Natalia Piórkowska, Malwina Brożyna, Tomasz Matys, Adam Junka

## Abstract

Invertebrate and larval model organisms such as *Drosophila melanogaster*, *Caenorhabditis elegans*, *Danio rerio* larvae, and *Galleria mellonella* are increasingly employed in biomedical, toxicological, and ecological research. Their behavioral responses serve as sensitive indicators of functional changes, yet traditional methods of observation remain low-throughput, subjective, and poorly scalable. Artificial intelligence (AI), including machine learning (ML) and deep learning (DL), has emerged as a powerful alternative, enabling automated and unbiased analysis of highly dimensional behavioral data. Here, we present the first systematic review comprehensively mapping the use of AI in behavioral analysis of invertebrate and larval organisms. Following PRISMA 2020 guidelines, we screened literature published between 2015 and May 2025. A total of 97 eligible studies were analyzed for model organisms investigated, AI methods applied, input data characteristics, preprocessing pipelines, model architectures, and evaluation metrics. We observed a steep increase in publications, from only 2 in 2015 to 97 by mid-2025, with the majority originating from the USA, China, and Germany. The most frequently studied organisms included *D. melanogaster*, *C. elegans*, and zebrafish larvae, alongside aquaculture and pest species. Since 2021, DL models, particularly convolutional neural networks (CNNs), including YOLO models, and pose estimation frameworks such as DeepLabCut have dominated the field, while supervised ML remains common for classification tasks, and unsupervised learning is primarily applied in exploratory clustering. Input data were typically video or image recordings, but reporting practices were highly inconsistent regarding resolution, frame rate, preprocessing steps, and model training details. Evaluation metrics also varied widely, limiting reproducibility and cross-study comparisons. To address these gaps, we propose a standardized reporting framework encompassing input data specifications, preprocessing pipelines, model architecture, and evaluation metrics. Such standardization will enhance transparency, reproducibility, and comparability across laboratories. AI-driven behavioral analysis has the potential to accelerate drug discovery, toxicology, and environmental monitoring while reducing reliance on vertebrate models in preclinical research.

## Introduction

Model organisms, particularly invertebrates such as insects (*Drosophila melanogaster*) and nematodes (*Caenorhabditis elegans*), have become frequently used tools in contemporary biomedical and pharmacological research. These model organisms retain a wide range of evolutionarily conserved biological pathways, including those governing neurodevelopment, innate immunity, stress response, and metabolism, enabling, to a reasonable extent, extrapolations to vertebrates, including humans. For example, *C. elegans* share homologs for up to 60–80% of human disease-related genes, underlining their utility in pharmacological and toxicological modeling *^1^*. In addition, their short life cycle, ease of handling, low maintenance costs, and minimal or no ethical constraints make them particularly suitable for high-throughput screening platforms*^2^*.

Larval stages of these invertebrates offer unique experimental advantages over adult forms, particularly in the context of neurobehavioral and toxicological assessment. Their structural and functional simplicity, reflected in reduced tissue and behavioral complexity, minimizes biological noise and facilitates more direct interpretation of experimental outcomes. Larval organisms often demonstrate high physiological sensitivity to chemical stimuli, making them ideal biosensors for detecting low-dose effects and early biomarkers of toxicity *^3^*. Metrics such as lethal dose 50 (LD₅₀), no observed effect concentration (NOEC), or changes in specific behavioral repertoires (e.g. exploratory behavior, locomotion, phototaxis) are frequently used to quantify toxic or pharmacological effects. For instance, *Danio rerio* larvae show quantifiable locomotor or startle responses to anxiolytics, antiepileptics, and neurotoxins *^4^*. *C. elegans* larvae have been instrumental in dissecting dopaminergic neurodegeneration and behavioral plasticity in response to diverse pharmacological compounds *^5^*. More recently, *Galleria mellonella* larvae have gained traction as a cost-effective alternative *in vivo* model in infection biology, immunotoxicology, and behavioral pharmacology. Their innate immune system shares several features with that of mammals, including conserved pathways for phagocytosis, production of reactive oxygen species (ROS), and synthesis of antimicrobial peptides (AMP). Unlike many invertebrate models, *G. mellonella* larvae can be maintained at 37°C *^6^*, which enables direct co-cultivation with human pathogens and exposure to human-derived biofluids or experimental agents. From a behavioral perspective, *G. mellonella* larvae exhibit distinct and measurable changes in posture, locomotion, and escape behavior under various stress conditions, including infection, inflammation, and chemical exposure, which can serve as functional readouts *^6,7^*. It positions *G. mellonella* as a promising model at the intersection of toxicology, translational medicine and behavioral neuroscience *^8^*.

Behavioral analysis has become one of the key approaches in preclinical and translational research, offering means of assessing the effects of pharmacological agents, neurotoxicants, and environmental stressors. Unlike morphological or molecular markers, which often capture static or endpoint changes, behavior reflects the dynamic output of interacting physiological systems, including the nervous, muscular, and endocrine networks *^9^*. As such, it can reveal functional alterations, long before structural damage becomes evident, making it a particularly valuable early indicator of adverse or therapeutic effects. In invertebrate models, behavior is increasingly employed to quantify neurophysiological integrity and systemic health. Parameters such as total distance traveled, swim velocity, angular displacement, turn frequency, freezing time, and burst activity are routinely extracted from these model organisms. For instance, such locomotor traces of zebrafish larvae are used to investigate their response to drugs targeting GABAergic (gamma-aminobutric acid-ergic), glutamatergic, or dopaminergic pathways *^10^*. The exposure to anxiolytics typically reduces thigmotaxis and increases center zone activity, while proconvulsants induce erratic movement or seizure-like bursting *^11^*. In *C. elegans*, chemotaxis assays, omega turns, and body bends are used to probe sensory integration and motor coordination. These behaviors are tightly regulated by conserved neurotransmitter systems and have been validated against known neuroactive compounds. Beyond classic dose–response studies, behavioral phenotyping is being leveraged in phenotypic screening pipelines and complex toxicity profiling. For instance, behavioral changes can serve as functional proxies for neurotoxicity, mitochondrial dysfunction, or developmental delay, especially when other biomarkers are inconclusive *^12^*.

The invertebrates’ behavior, especially in larval form, is also highly responsive to environmental toxins, food additives, and biological fluids, making it applicable in ecotoxicology, forensics, and environmental monitoring. Functional endpoints like latency to stimulus response or recovery time after exposure have been proposed as alternative metrics for evaluating sublethal effects in regulatory toxicology frameworks *^13^*. Importantly, behavioral endpoints offer translational relevance. Phenotypes such as hypoactivity, hyperactivity, ataxia, startle failure, or habituation deficits often mirror clinical symptoms in neurological and psychiatric disorders. As such, larval behavior may be perceived not just as a readout of gross toxicity, but as a correlate of cognitive, emotional, or neuromotor function. Moreover, because behavior can be continuously and non-invasively monitored, it is well-suited for longitudinal studies, dynamic dose modeling, and real-time phenotypic screening. In summary, behavioral analysis in invertebrate and larval organisms may bridge the gap between molecular changes and organism-level function, enabling high-sensitivity evaluation of pharmacological and toxicological effects in a scalable manner.

Traditionally, behavioral analysis relied on two main mechanisms. Either behavior was observed by a qualified researcher, or an assay was designed to measure a specific aspect of behavior *^14^*. For instance, human-based scoring of number of courtship behaviors in genetically mutated fruit flies showed that single gene splicing can specify complex innate behavior *^15^*. Nevertheless, such traditional approaches are time-consuming, low-throughput and prone to observer bias. Moreover, they are particularly inadequate for detecting subtle or transient phenotypes, especially those involving complex temporal dynamics or multi-dimensional movement patterns *^16^*. Semi-automated tracking systems have improved the throughput and objectivity of behavioral studies but still depend heavily on human input. These systems typically extract positional data (e.g. centroid tracking, velocity, trajectory curvature) from video recordings, and provide derived metrics such as distance traveled, acceleration, or turning angles. For instance, locomotion on *D. rerio* in larval stage, treated with early-stage therapeutics, was validated as an early-stage toxicity screen *^17^*. Undoubtedly, being an improvement on traditional behavioral analysis, these methods often struggle with occlusions, low contrast, complex postures or with overlapping individuals in group settings. Moreover, predefined rule-based classifications (e.g. “freezing” if speed reduces to a predefined value) may oversimplify complex and/or ambiguous behaviors. Importantly, both manual and semi-automated methods lack, to the major extent, scalability and adaptability *^18^*. Behavioral experiments generate large volumes of video data, yet traditional tools are ill-equipped to handle these datasets at scale.

Building upon traditional and semi-automated approaches, recent advances in artificial intelligence (AI) have begun to transform behavioral analysis landscape. Ranging from organisms’ automatic detection, classification, thorough pose estimation, tracking and behavior clustering AI-based techniques revolutionized the field of behavioral biology. Machine learning (ML), which is a core branch of AI, deals with programming computers to “learn” patterns in data. It can be done by providing labeled data (e.g. images) to the algorithm (referred to as supervised learning (SL)) or by automatically grouping data into clusters of alike characteristics without training the model on predefined labeled data (referred to as unsupervised learning (UL)). These models are relatively straight-forward to build, work well on smaller, preselected datasets and in general are easily interpretable. On the other hand, they still rely on human input at the model training stage, selecting which behavioral parameters should be considered by the model-called feature selection. A dynamically growing group of deep learning (DL) approaches aims at tackling some of the limitations of classical ML. This group includes the most popular convolutional neural networks (CNN), recurrent neural networks (RNN) and Transformers, among others. These models take a more flexible and objective approach to feature selection and work on highly dimensional and unstructured data. Unfortunately, this comes at the cost of the requirement for a significantly larger training dataset and less transparent and reproducible methodology.

Building upon this emerging potential, it becomes essential to delineate the concrete tools that operationalize these advances for experimental biology. A growing ecosystem of open-source platforms now provides biologists with accessible solutions for pose estimation, multi-animal tracking, and behavior classification. While originally developed within computational sciences, many of these frameworks have been adapted to laboratory contexts, lowering the technical threshold for their use. Table 1 introduces the most prominent of these tools, such as DeepLabCut, SLEAP, and JAABA presenting their application in behavioral analysis.

**Table 1.** Summary of the key AI tools developed for or used in behavioral analysis of invertebrate and larval model organisms.

| AI tool | Description | Precursors |
| --- | --- | --- |
| DeepLabCut | Open-source software package for markerless 2D and 3D pose estimation of animals. Based on transfer learning with CNN model achieves excellent results with little training data. | none |
| SLEAP | Open-source package for general-purpose, multi-animal pose tracking. CNN model with pretrained wights, but can also enable transfer learning. | LEAP |
| FLLIT | FLLIT stands for Feature Learning Leg Segmentation and Tracking. It is a graphical user interface (GUI)-enabled program, MATLAB compatible. Analyzes gait parameters (incl. lurching steps, short strides). | none |
| WormPose | Open-source package. CNN-based model for estimating challenging poses of *C. elegans*, in 2D. Generates synthetic worm-real worm pair for training. | none |
| StrongSORT | Multi-object tracking algorithm. | DeepSORT |
| Z-LAP Tracker | Zebrafish Larvae Position Tracker. Modification of DeepLabCut, specifically trained for zebrafish larvae. Quantifies behavioral parameters. | none |
| JAABA | Open-source program. Annotate and classify specific behaviors based on kinetic features. | none |
| Stytra | Open-source tool for tracking, visual simulations and close-up experiments. It is a graphical user interface (GUI)-enabled software shown to work in conjunction with e.g. DeepLabCut in behavioral experiment. | none |
| Anipose | Open-source python toolkit for 3D markerless pose estimation, built on 2D pose estimation tool DeepLabCut. | none |

These AI tools, along with the fast-evolving field of AI-enabled analysis undoubtedly transformed how researchers study behavior. This interdisciplinary field of biology brings together expertise from computer science, neuroscience, microbiology to ecology, enabling sophisticated approaches to capture the non-linear dynamics and complexities of behavior that were previously difficult to quantify or even detect. Despite the growing interest and potential, a survey of the literature reveals that, out of eight identified reviews *^19–26^*, none comprehensively maps the current landscape and critically evaluates the methodological advancements. Consequently, the field suffers from fragmented information, with no clear guidelines for applying these tools across diverse invertebrate and larval models. This systematic review aims to fill that gap by summarizing the state of AI-based behavioral analysis in these organisms, highlighting both the limitations of current methodologies and the opportunities they present, and providing a forward-looking perspective on their potential translational impact in biological research.

## Methods

This systematic review was conducted in accordance with the Preferred Reporting Items for Systematic Reviews and Meta-Analyses (PRISMA) 2020 guidelines *^27^*. Its aim was to identify recently published scientific literature on AI methods used in behavioral analysis of invertebrate and/or larval model organisms, summarize current trends in research and propose future directions. The following subsections present detailed methodology.

### Search strategy

In order to conduct thorough and systematic search of scientific literature, the Sample, Phenomenon of Interest, Design, Evaluation, Research type (SPIDER) tool was implemented *^28^*. Information summarized in Table 2 below aided in formulating database search queries (Table S1) and inclusion/exclusion criteria (Table 3).

**Table 2.** The overview of the SPIDER tool used to formulate search queries for the systematic review.

|  |  |
| --- | --- |
| **Sample** | Invertebrate and larval model organisms |
| **Phenomenon of Interest** | Use of AI methods for behavioral analysis, including detection. |
| **Design** | Published literature, including preprints, that included sufficient methodology to understand and evaluate AI models. Studies where behavioral data was acquired via video, image or motion tracking systems. |
| **Evaluation** | Eligible studies report the following: organism model investigated, type of AI-model used, type and quantity of input data, source of input data, preprocessing and analysis steps, where applicable model architecture, and model evaluation metrics. |
| **Research Type** | Quantitative, exploratory, and comparative studies. |

**Table 3.** Inclusion/exclusion criteria applied during the search procedure in the systematic review.

| Inclusion criteria | Exclusion criteria |
| --- | --- |
| **Stage 1: Initial screening of title and abstracts** |  |
| Both empirical studies and literature reviews that used AI-based methods |  |
| Studies investigating behavioral analysis including detection, movement, activity patterns, social interactions |  |
| Studies investigating invertebrate and/or organisms in larval stage | Studies focusing entirely on adult, vertebrate organisms |
| Studies where organisms and their behavior was recorded as images, videos or motion trajectories |  |
| Published in the period 01.01.2015-31.05.2025 |  |
| Full text available in English |  |
| Preprints | Publication type including bachelor's, master's and doctorate thesis and book chapters |
| **Stage 2: Full text analysis** |  |
|  | Studies using only traditional analytical methods to study behavior in model organisms |
|  | Studies recording behavioral analysis via a microscope. Authors define microscope as a static device using series of magnifying lenses to obtain an image or video in a macro scale |
|  | Studies with insufficiently described methodology and results, making it impossible to answer the stated RQs |

Furthermore, the following Research Questions (RQ) were identified and used to guide the feature extraction and analysis stages, ensuring comprehensive and detailed overview.

RQ1: What is the current landscape of AI-based research in behavioral analysis of invertebrate and/or larval organisms? This question investigates model organisms used, types of AI models, their applications and overall scientific interest in the field.

RQ2: What types of input data and behavioral features are used in these models? This question aims to assess how behavior is recorded and quantified.

RQ3: How effective and readable are AI-based models and their analysis pipelines? This question explores preprocessing steps, model architecture (where applicable), and evaluation metrics.

### Search procedure

The search was completed on 31^st^ May 2025, and included literature published between 1^st^ January 2015 and 31^st^ May 2025. This time period was chosen to accurately represent the current landscape of the dynamically developing field.

PubMed, Scopus, Web of Science, IEEE Xplore, and ACM Digital Library databases were searched, as per defined queries (Table S1). Additionally, to saturate the search, the first 200 publications from Google Scholar search engine were added. The literature was aggregated in Mendeley.

The inclusion criteria were tailored to identify research that investigated behavioral analysis using computational approaches, including image processing pipelines, simple ML algorithms, DL methods, and other AI-based approaches. The studies had to focus, or at least test algorithms on invertebrate organisms or organisms in larval stage. Moreover, to broaden the scope of analyzed publications, authors decided to include papers which used AI models to investigate behavior (including movement, activity patterns or social interactions) or perform organism’s detection. The organism had to be recorded via a camera or motion sensor, and not with a microscope. This approach ensured that recording sources focused on whole organisms, rather than sub-parts or cells, and downstream analysis of methods could be compared across research. Finally, both peer-reviewed literature and preprints, published between 1^st^ January 2015-31^st^ May 2025 were included.

The main exclusion criteria included analyzed behavior using only traditional analytical tools or did not analyze behavior at all. Studies that focused exclusively on adult vertebrates were also excluded. Finally, studies that met inclusion criteria, but did not contain sufficient methodological details on model used, its evaluation metrics, source, type and quantity of input data were also excluded. This ensured both that only studies of high quality were included, and that authors could compare models and their characteristics across studies.

The initial database search yielded 1745 results. Out of these records, using Mendeley, 471 duplicates were identified, manually verified and removed. Next, in the first stage of selection, authors screened papers based on titles and abstracts. At this stage, 1123 records were removed. The remaining 151 publications underwent thorough analysis and based on full text 54 of these were further excluded. The literature search was done independently by two authors and, where necessary, discrepancies were resolved through discussion and consultation with the rest of the authors. Below, the PRISMA diagram *^27^* summarizes the search procedure (Fig 1).

**Figure 1.**
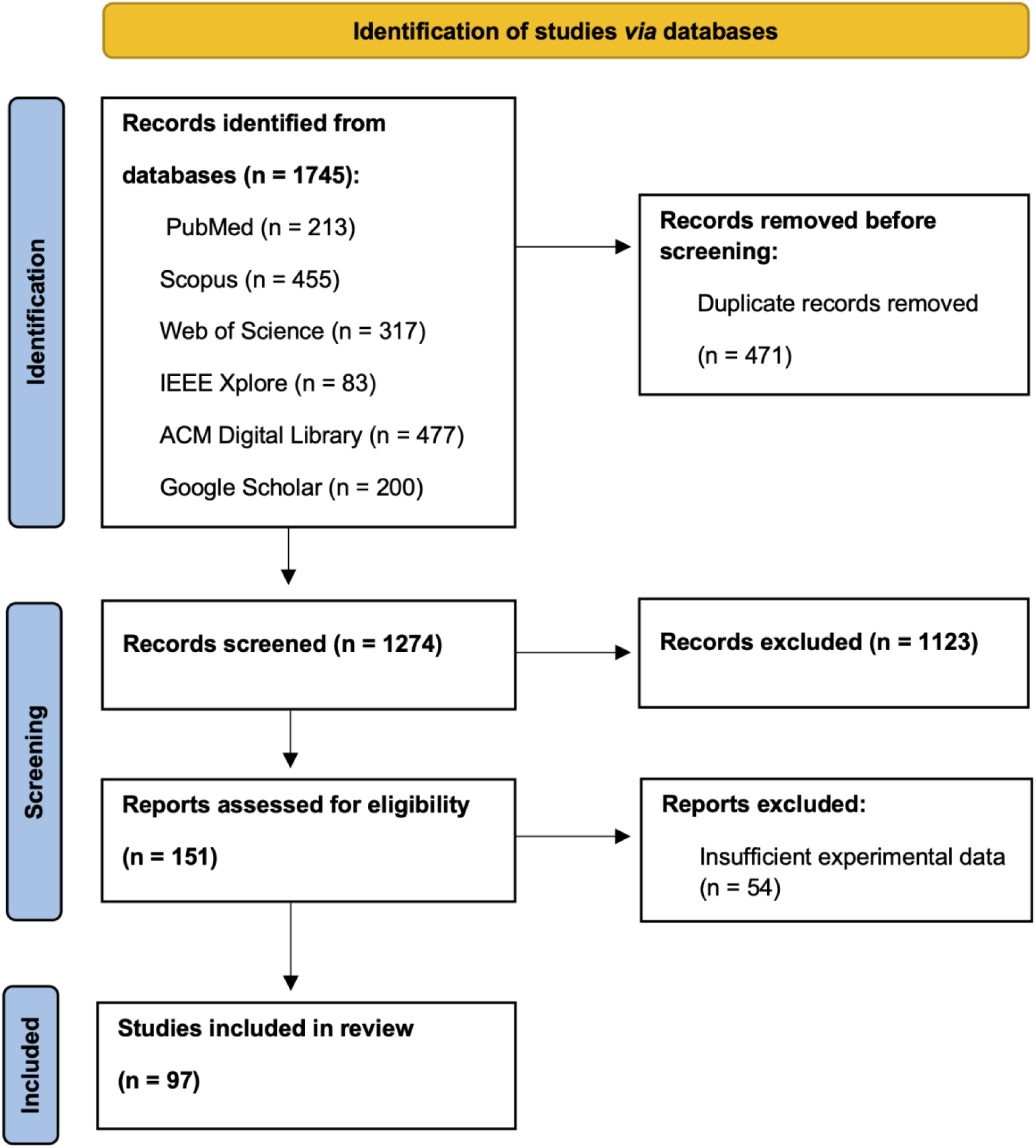
The PRISMA diagram showing systematic search procedure.

### Features extraction

The literature included in the final analysis underwent a detailed feature extraction process. Authors decided on a set of key information, relevant to comprehensively answer previously stated research questions. These include:

1. publication details, publishing journal, publication year, model organism investigated, AI model used, model’s task and aim of the analysis;
2. type and quantity of input data, source of input data, behavioral characteristics of organism analyzed;
3. preprocessing steps, model architecture (where applicable), evaluation metrics reported.

Data extraction was carried out by two authors and cross-checked to mitigate extraction errors. The full set containing all extracted data is available in supplementary materials (Table S2-4).

### Data analysis

Aggregated results were summarized in figures and supported by descriptive statistics were applicable. Data was aggregated in Excel, and preprocessed and visualized using RStudio (R version 4.5.0). The results were structured around research questions and authors formulated qualitative assessment of methodological rigor.

When studies investigated multiple model organisms, employed hybrid models (meaning more than one model was used) or different types of input data, each instance was indexed and reported separately. Exemption to the rule were studies which compared multiple models. In those cases, only the model evaluated as the best performer was reported. As a result, totals for some statistical comparisons may exceed the total number of analyzed studies. For variables like a country of study and research area investigated, only the primary/main instance was considered. Finally, for variables such as preprocessing steps, model architectures, and evaluation metrics, key information was summarized, with data grouped and reported selectively to highlight elements most relevant to the analysis. Authors decided on this approach to present extracted results in the most informative format, providing a comprehensive landscape of AI methods utilized in behavioral analysis of invertebrate and larval model organisms.

## Results

### Temporal and geographical trends

Analysis of 97 publications, included to review, showed a growing interest in the field of AI-based research on behavior of invertebrate and larval model organisms. Starting from only 2 papers being published in 2015*^29,30^*, to 43 papers (34 in peer-reviewed journals *^29–62^* and 9 in preprints and conference proceedings *^63–71^*) by 2021, and already 8 papers published in 2025 *^72–79^* (Fig 2, Table S2). Notably, 43% of literature originated from the USA, China and Germany (Fig 3b), with 80% *^30,42,55,76,79–86^*, 60%*^35,72,73,75,78,87^* and 100%*^37,40,43,47,49,62,88^*, respectively being published in journals scored with impact factor 3 and above (Fig 4). This concentration of high-quality research output reflects the leading roles of these countries in both AI and life sciences fields *^89,90^*. Countries in South America, Asia and Eastern Europe also undergo research in the field (Fig 3a, Fig 4), indicating global engagement possibly aided by widely accessible computational power for model development. Interestingly, the majority of published research in the last decade appeared in journals with Impact Factor greater than 3 (74%) *^30,34–37,39–50,53–62,72–88,91–100^*(Fig 4). This places AI research on behavioral analysis of invertebrates and larval organisms as a significant, trending topic in the sub-field of science.

**Figure 2.**
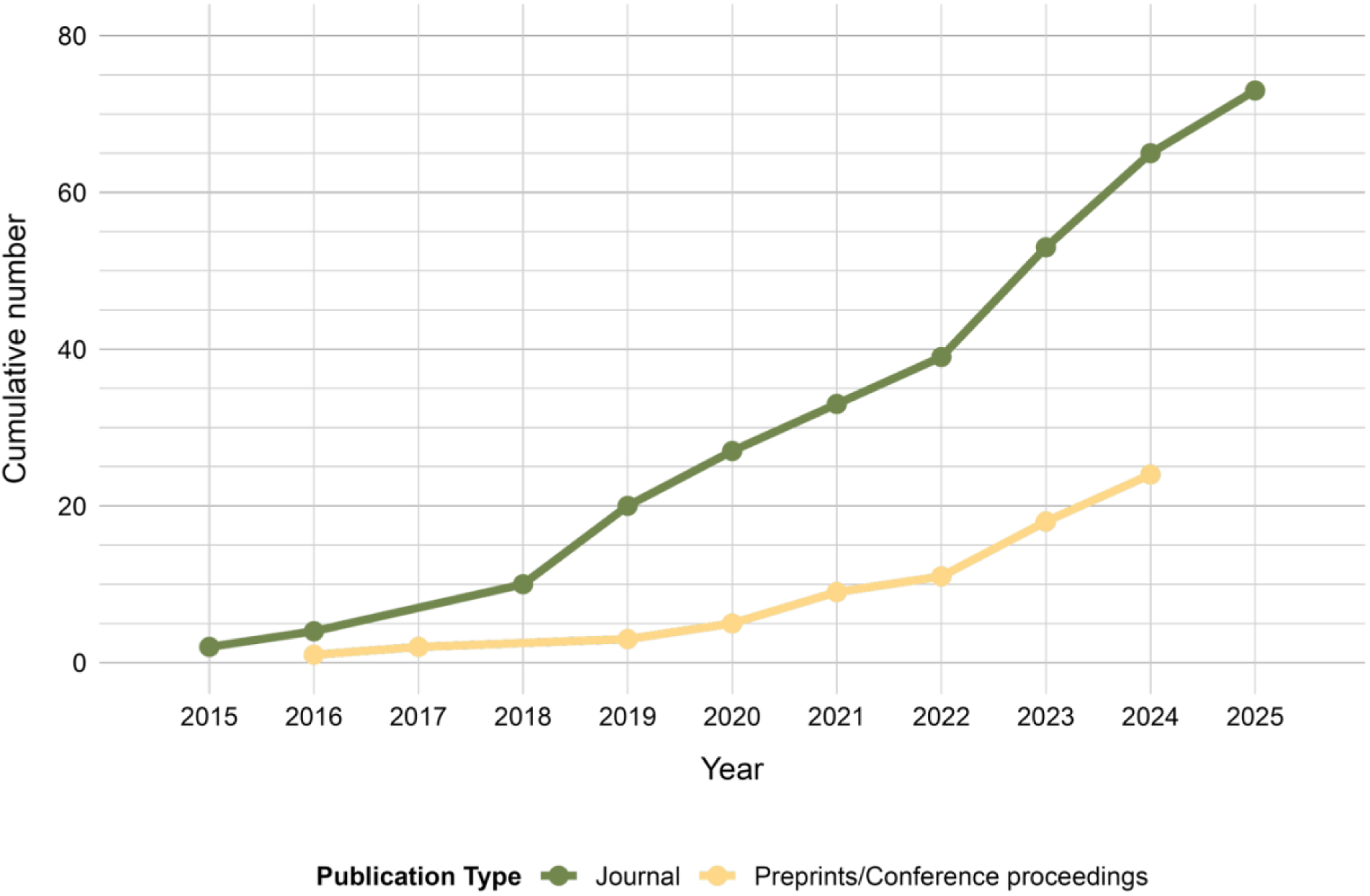
Cumulative number of research published between 2015-2025.

**Figure 3.**
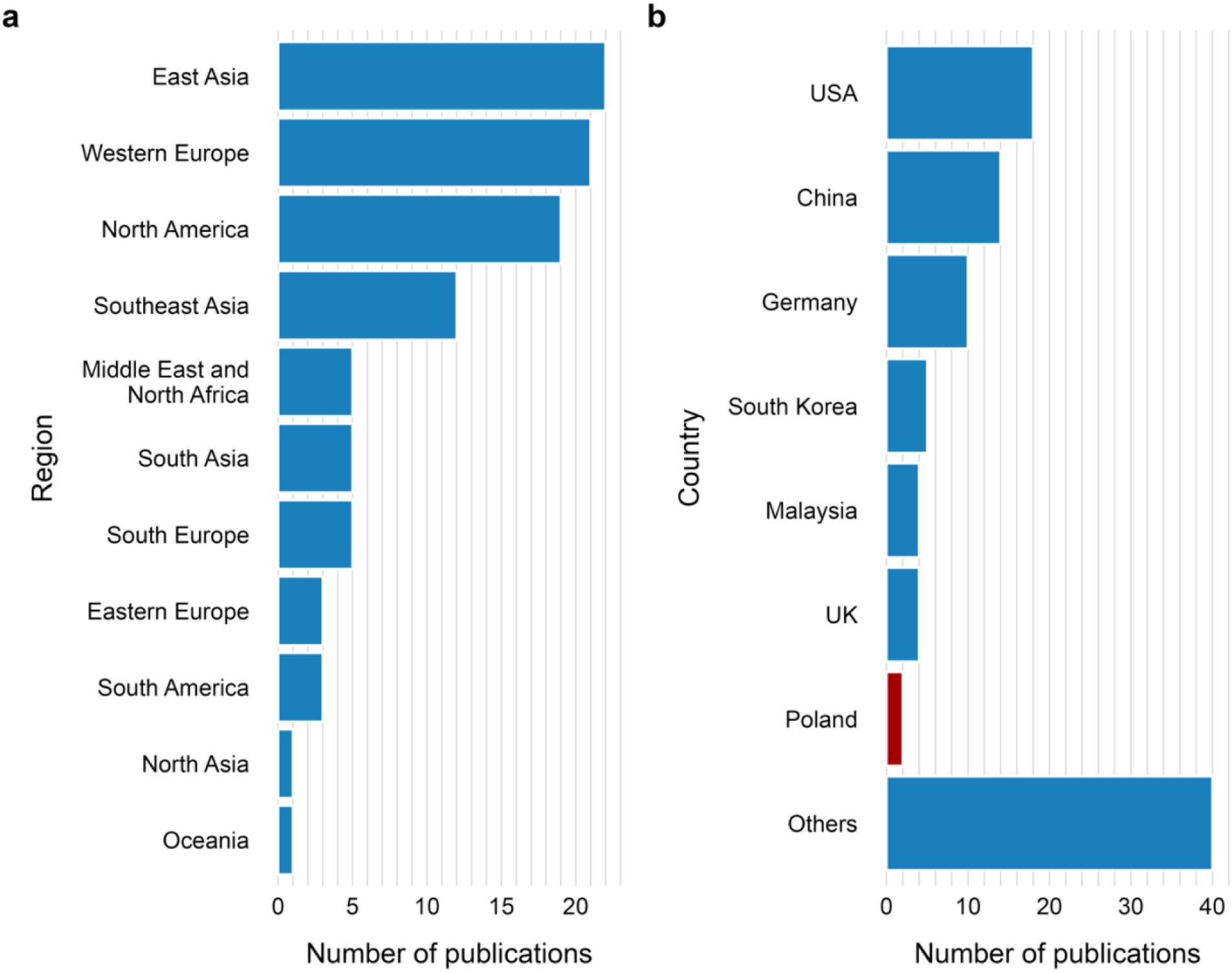
Number of peer-reviewed articles, preprints and conference proceedings published between 2015-2025, by geographical region (a), and by country (b). The country of authors’ origin is red-colored.

**Figure 4.**
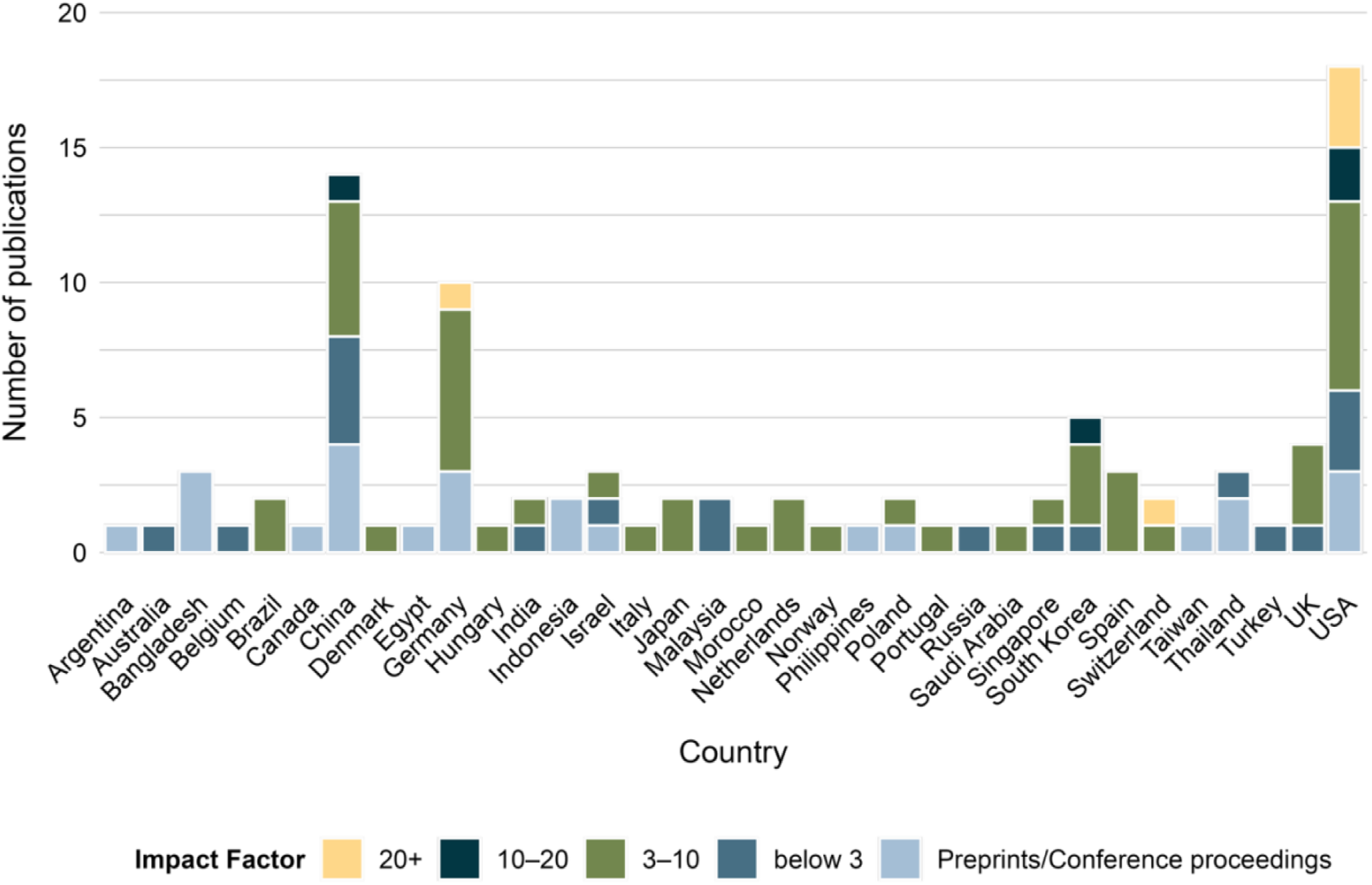
Number of articles published between 2015-2025 by country, grouped by journal’s Impact Factor.

### Model organisms investigated

Over 50% of published research utilized widely recognized model organisms in biology, including D. melanogaster (22%) *^30,37,41,42,44,46,47,51,53,55,58,62,63,71–73,79,81,84,86,88,101–103^*, C. elegans (18%)*^32,33,36,43,45,50,57,59,72,78,80,87,91,92,94,100,104–106^* and D. rerio, in larval form (11%)*^29,34,40,42,49,72,82,83,85,107–109^* (Fig 5, Table S2). Within this group of organisms, research was conducted on larval forms in approximately 38%*^29,34,40,42,49,53,62,72,80,82,83,85,87,88,104,107–109^* (Table S2). These organisms were chosen as model organisms across almost all identified research fields (Fig 6), including development of automation tools for ethology, genetics, ethology, agriculture and farming, neurobiology, aging, environmental toxicology, pharmacology and toxicology. Other significant groups of model organisms investigated were shrimps (incl. *Litopenaeus vannamei*)*^64,68,75,76,110–113^* and mosquitos (incl. *Culex annulirostris, Aedes aegypti*)*^38,52,69,114–117^*, mainly investigated in aquaculture and epidemiology, respectively (Fig 6, Table S2). Moreover, various pests, moths, fish larva and insect species found applications in agricultural and farming*^35,48,54,56,60,67,70,74,93,118–124^*, aquacultural*^65,95,96,98^*, ethological*^31,125,126^* and ecological studies (Fig 6).

**Figure 5.**
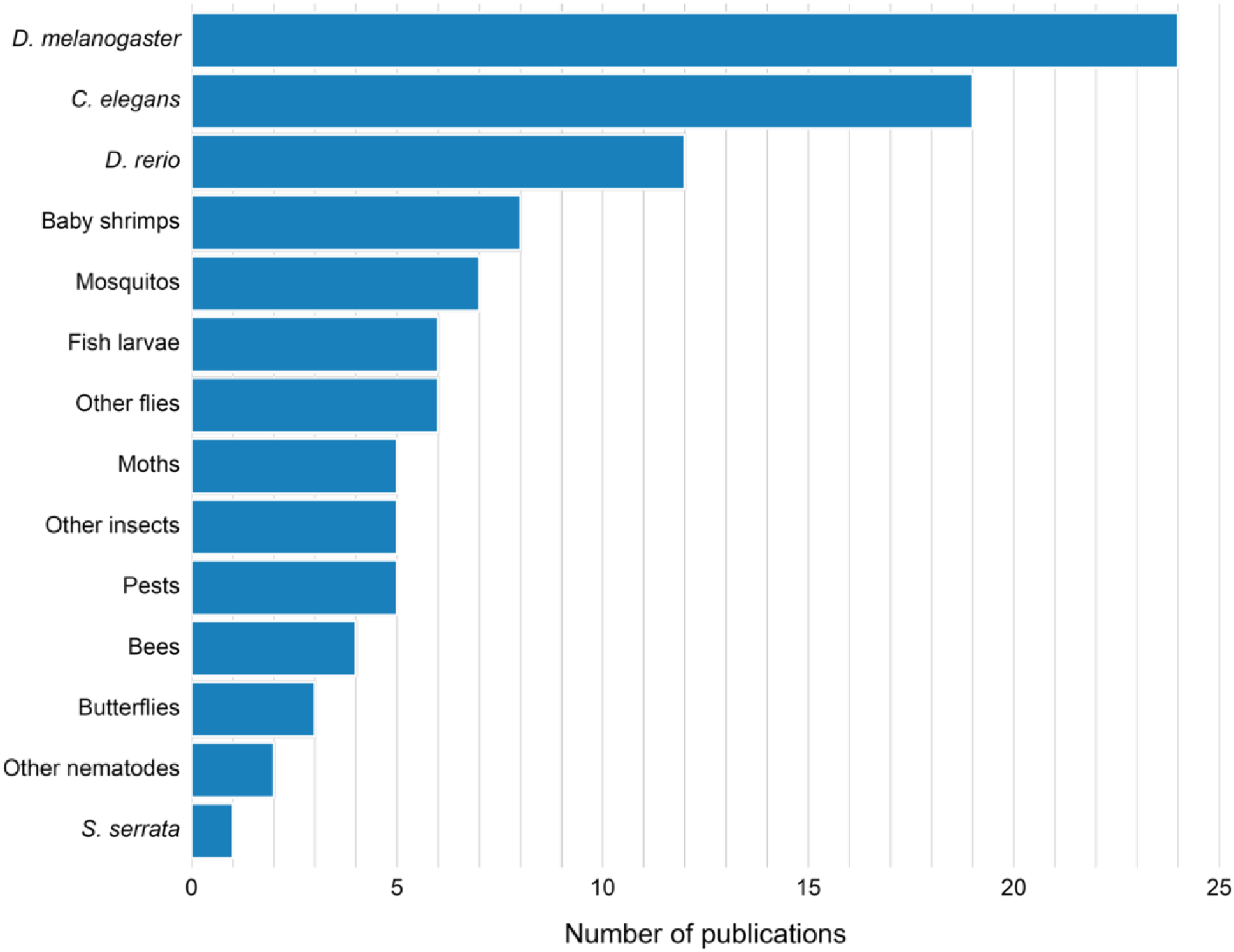
Model organisms investigated in analyzed research papers.

**Figure 6.**
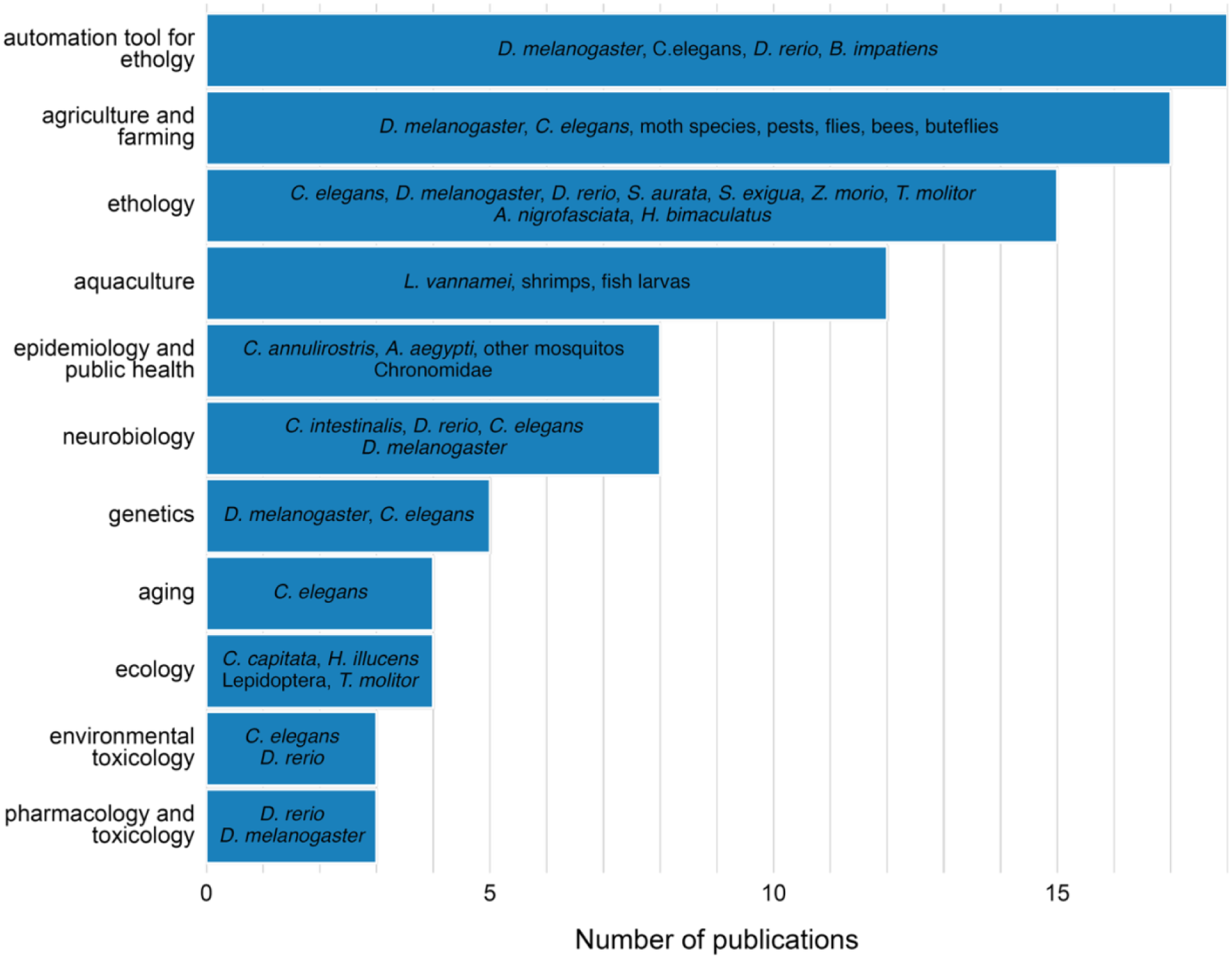
Investigated model organisms, according to the research field.

### Overview of models used in behavioral research

Deep learning was the dominant method used in the field of AI-based behavioral analysis of invertebrate and larval organisms, followed by supervised learning and unsupervised learning (Fig 7a). The less frequently chosen model group was image analysis, defined as semi-automatic approach consisting of the pipeline of code that processes input data without the element of learning (Fig 7a). Some of the examples of algorithms used in this group of models include Adaptive Shape Model *^88^*, or adaptive thresholding incorporated into analysis pipeline *^33^*. The least popular method, appearing only three times across investigated period, was the statistical approach (e.g. Particle filtering or Reversible Jump Markov Chain Monte Carlo algorithm)*^53,62,127^*. Until 2020 the development of AI-based methods was balanced between the four main groups: DL, supervised learning, unsupervised learning and image analysis algorithms, appearing in 12 (29%)*^30,37,38,41,42,45–48,53,54,66^*, 11 (27%)*^31,35,36,39,43,44,51,52,65,71^*, 9 (22%)*^29,32,34,39,45,49,63^* and 7 (17%)*^33,40,50,63–65,71^* papers, respectively (Fig 7a). After 2020, however, the number of papers focusing on developing DL algorithms skyrocketed to the total of 74 (62%)*^30,37,38,41,42,45–48,53–61,66–70,72,73,75,78–87,91–101,103–127^* by 2025, while supervised and unsupervised learning approaches grew in popularity at a steady rate, adding up to 20 (17%)*^31,35,36,39,43,44,51,52,65,71,74,76,79,84,85,97,102,104,108^* and 13 (11%) *^29,32,34,39,45,49,63,79,83–85^*, respectively (Fig 7a). The drastic increase in development of DL approaches can be attributed, to some degree, to the development of CNN-based markerless pose estimation tools called DeepLabCut *^37^* and SLEAP *^81^*, which became widely used tools in behavioral analysis and precursors for other pose estimation algorithms (e.g. Z-LAP tracker *^83^*).

**Figure 7.**
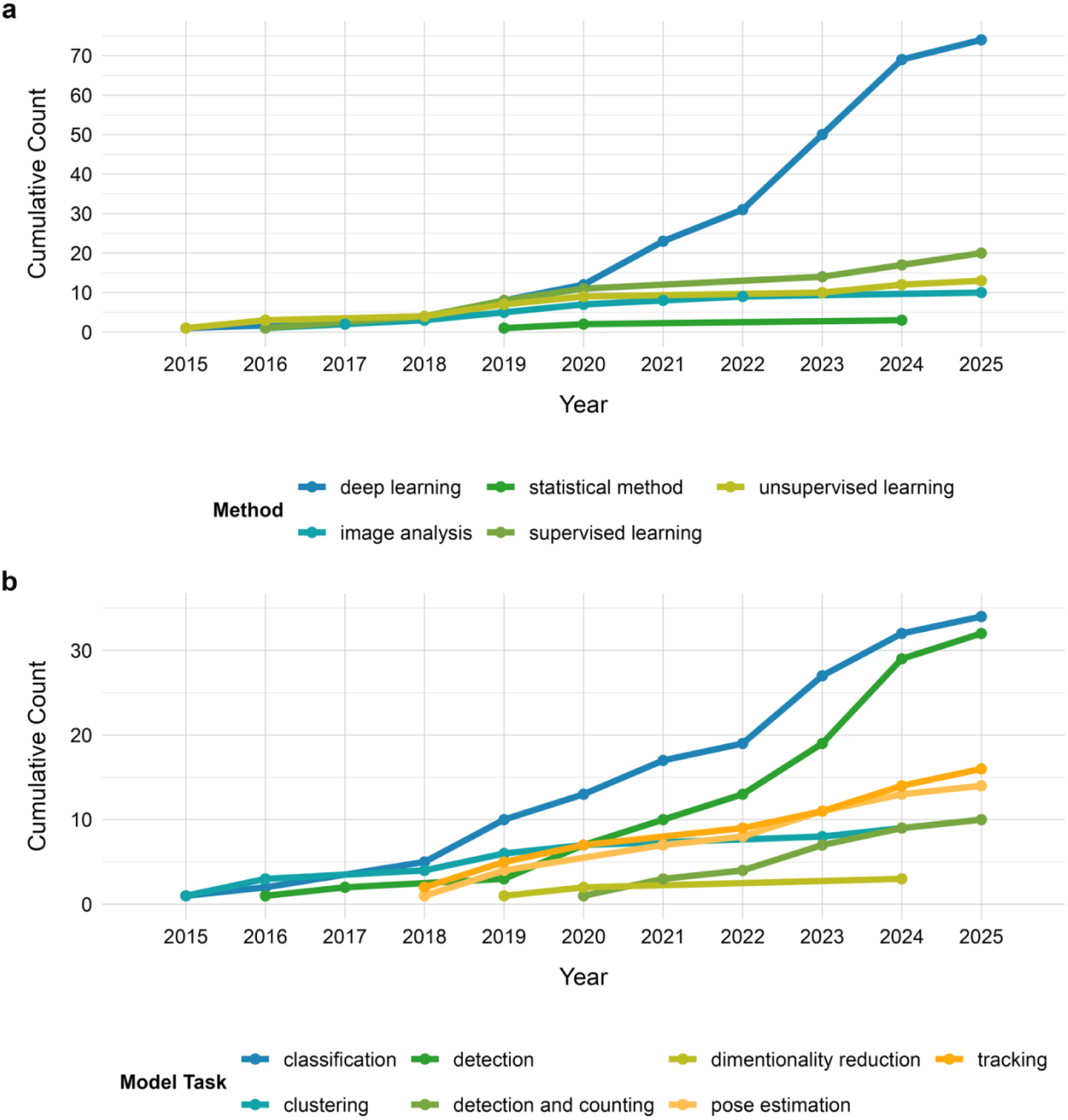
Overview of AI-based approaches in behavioral analysis of invertebrate and larval model organisms. Grouped by method (a) and model task (b).

Investigating types of tasks models performed, the dominant ones are classification*^30,31,35,36,38,39,43,45,46,52,54,57,61,65,67,69,71,74,79,84,85,91,92,97,100,102,104,107,108,113–115,126^*and detection*^48,50,53,63–65,68,70,75,77,78,87,88,92,93,95,98,99,101,103,106,111,116,118–123,125,127^* (34 and 32 uses, respectively), followed by tracking*^33,36,40,44,51,62,71,73,79,80,86,104,105,109,125,127^* and pose estimation*^37,41,42,47,55,58,59,72,81–84,97,108^* (16 and 14, respectively), clustering*^29,32,34,39,45,49,63,79,83,85^* and detection and counting*^56,60,66,76,94,96,110,112,117,124^* (10 uses, each), and less common dimensionality reduction techniques*^45,49,84^* (3 uses) (Fig 7b).

Looking at the models’ profiles, grouped by method and type of task performed (Fig 8), DL was used in almost all classes of tasks, as an exclusive approach for pose estimation as well as the dominant approach in detection and detection and counting tasks. DL and supervised learning methods were used in similar relative frequency in classification tasks. Tracking tasks used almost all methods, while clustering, and dimensionality reduction tasks, as an inherently separate group of tasks, used unsupervised learning approaches.

**Figure 8.**
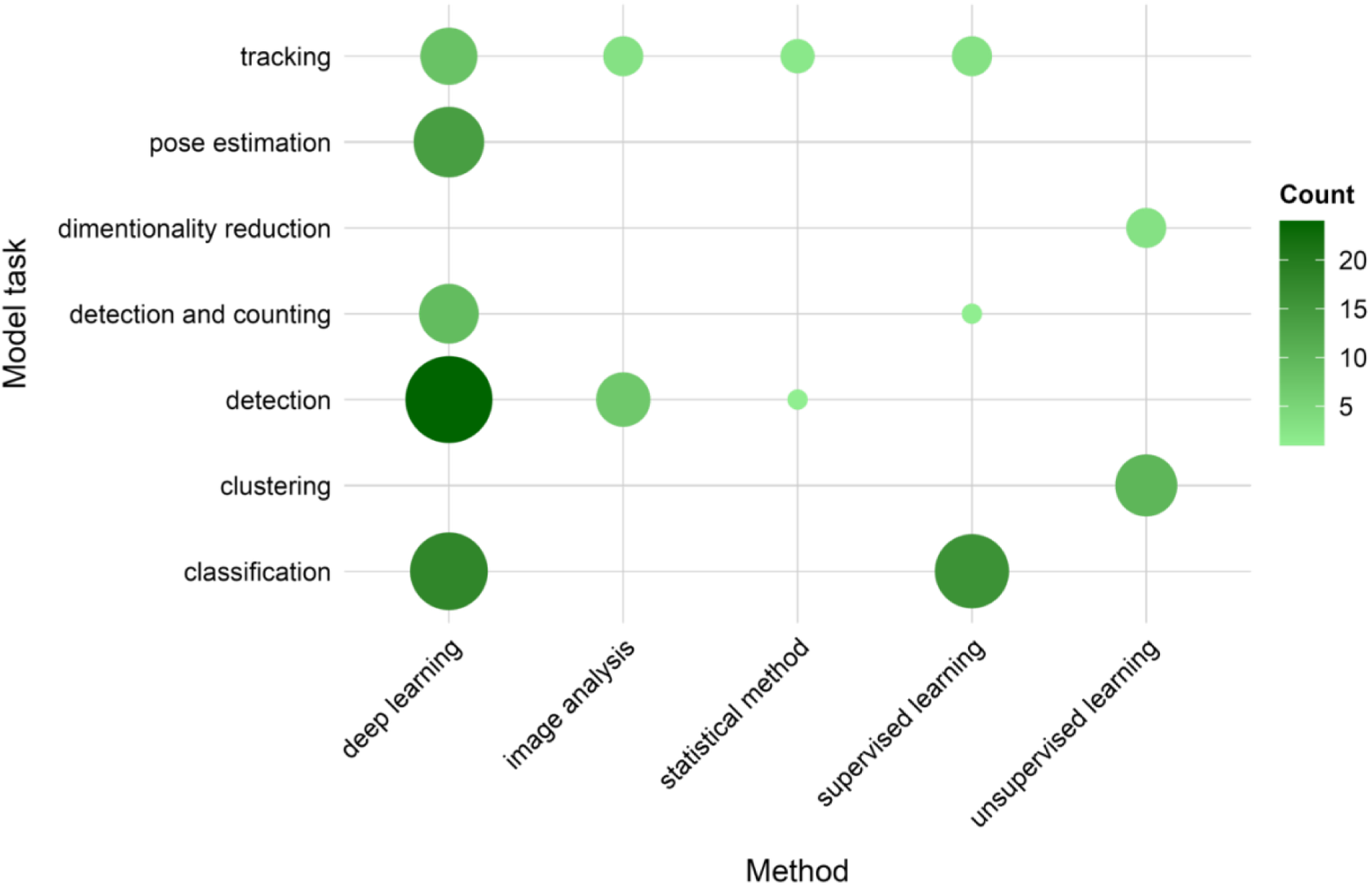
Distribution of AI models’ tasks used in different types of methods.

Diving deeper into the specific algorithms, within a subgroup of DL approaches the most common algorithms used were based on CNN architecture, and included YOLO (You Only Look Once) algorithms*^68,70,87,99,101,103,105,106,109,120,122,123,125^* (13 uses), Faster R-CNN*^60,80,93,94,96,121,124^* (7 uses), markerless pose estimation tool DeepLabCut*^37,55,84,97,108^* (5 uses), along with other pose estimation tools that either were directly based on DeepLabCut (Z-Lap Tracker*^83^*) or employed a similar algorithm (LEAP*^42^*, SLEEP*^81^* and WormPose*^59^*). A less common, yet increasingly developed DL methods that appeared in published research were transformers and RNN architectures. While transformers (RT-DETR, VITPose and KAN Transformer) were used in variety of tasks (detection*^98,119^*, pose estimation *^72^* and tracking*^73^*, respectively), RNN models (LSTM) were used twice for classification of *C. elegans* swimming patterns *^91,100^* and once for predicting larval movement trajectory 1.5 seconds into the future *^125^*.

Taking a closer look at the supervised, or classical ML approach, the three most commonly used algorithms included Support Vector Machines (SVM), Random Forest (RF) and K-nearest neighbors (KNN), being used as a stand-alone classification algorithm or followed by simple image analysis (4*^31,35,43,52^*, 2*^74,102^* and 1*^65^* use, respectively), in conjunction with pose estimation tool to classify poses into investigated phenotypes (2*^79,108^*, 3*^97,108^* and 1*^97^* use, respectively) and less commonly as an initial classifier to reduce dimensionality for deep learning (RF and KNN followed by Siamese Neural Networks Twin-NN an Twin-DN *^85^*), or following unsupervised clustering into behaviorally distinct groups (agglomerative clustering algorithm followed by KNN*^39^*).

The unsupervised approach was most commonly applied in clustering algorithms *^29,32,34,39,45,49,63,83^* (e.g. k-means clustering, MotionMapper, n=8) and less commonly in dimensionality reduction *^49,84,85^* (e.g. UMAP, SOM, n=3). The clustering method was used to group together animal poses*^79,83,84^*, as a stand-alone tool to group behavioral patterns such as swimming or movement patterns*^29,32,34^*, and in conjunction with other approaches to cluster behavior as a preparation step for further ML or DL analysis *^39,45^*.

Finally, image analysis approach, was documented to be used until 2021 in 8 papers (Fig 7), as a stand-alone tool for detection or tracking*^33,40,50,64,88^*, as a preprocessing step for classical supervised learning*^65,71^* and as a detection technique for clustered animals*^39^*. It is possible that with the development of DL approaches and higher complexity of behavioral research, image analysis was less frequently reported in literature’s methodology or even neglected in favor of presenting end-to-end DL pipelines.

Of note, several DL models (CNN-based models incl. YOLO and Faster R-CNN) had been pretrained on COCO or ImageNet datasets for detection and classification tasks*^77,78,80,84,91,93,99,101,103,105,114,115,121^*. This procedure accelerated training, allowing for a smaller number of input data required to achieve high accuracy results. However, it can be seen from model’s task profile, pretraining had only been advantageous for relatively simple detection tasks. On the other hand, transfer learning had been used in a wider, and more computationally demanding group of tasks, such as CNN-based algorithms for classification and pose estimation*^37,38,55,56,113,127^*. Finally, 2 papers utilized active learning for pose estimation tasks *^41,42^* allowing for further training in real-time without user intervention. One of the examples of classification into abnormal and normal swimming patterns used semi-supervised approach *^107^*, meaning the model was trained on control data only and was able to classify abnormal behavior as a distinct pattern from learnt control dataset.

### Overview of input data

The vast majority of experimental data for model training came from in-lab experiments with only 14% of papers training models on public datasets*^36,46,54,62,78,88,92,100,105,110,114,116,119,126^*. The most popular data acquisition technique was image (RGB, monochrome or greyscale) or video capture using a single camera (Table S3). Less frequently researchers used:

- Multiple synchronized camera set-up (DL models for pose estimation*^41,55,58^*, tracking*^86,127^* and classification*^30^* tasks)
- Specialized cameras such as hyperspectral*^35,74,99^* or CCD cameras*^45,53,91^*
- Cartesian multi-view robot (DL model for detection and counting*^94^*)
- Specialized laboratory incubators with tracking systems (e.g., ZebraBox*^107^*, FlyBowl*^71^*, WormTracker*^32^*, Ethovision XT*^102,107^*).

Interestingly, 11 papers used high-speed cameras to capture animal behavior*^29,31,34,40,44,49,51,65,73,79,108^* while 3 used IR-sensitive cameras*^34,39,40^*. In 10 experiments data was acquired via a smartphone camera*^43,52,68,95,96,101,109,112,113,123^*. Finally, 5 papers generated synthetic data*^59,66,77,82,94^*, on top of in-lab acquired material.

Unfortunately, the level of detail on source of input data was not reported cohesively across the literature. While some papers reported detailed specifications on input data (e.g. resolution, recording speed in a form of frames per second (fps), shutter speed) or camera used (e.g. model, filters, lenses), others did not report any details, even as basic as camera model (Table S3).

This heterogeneity in reported data is also visible in the format of reporting input data used to train the model. As seen on Fig 9a authors reported quantity of input data in the units of images*^33,35,37–40,42,44–47,51,54,56,58–60,63–65,68–70,72–74,77,83,84,88,91,93,94,98–101,105–107,109,112,113,115,116,118,119,121,123,124,126,127^*(54%), annotated organisms*^30,41,43,53,55,61,67,75,76,78–81,87,92,95–97,103,104,110,111,114,120,122,125^* (27%), videos*^31,32,36,52,57,82,108,117^* (8%), poses and swim bouts (specific type of pose especially for *D. rerio* larva)*^29,34,49,50,62,71,86^* (7%) and least commonly motion trajectories*^32,85,102,107^* (4%). Moreover, as seen in Fig 9b, the actual number of input data ranges from 100-1,000,000 in all input data categories, from less than 100 input data (in a form of videos) to over 1,000,000 input data (in a form of images).

**Figure 9.**
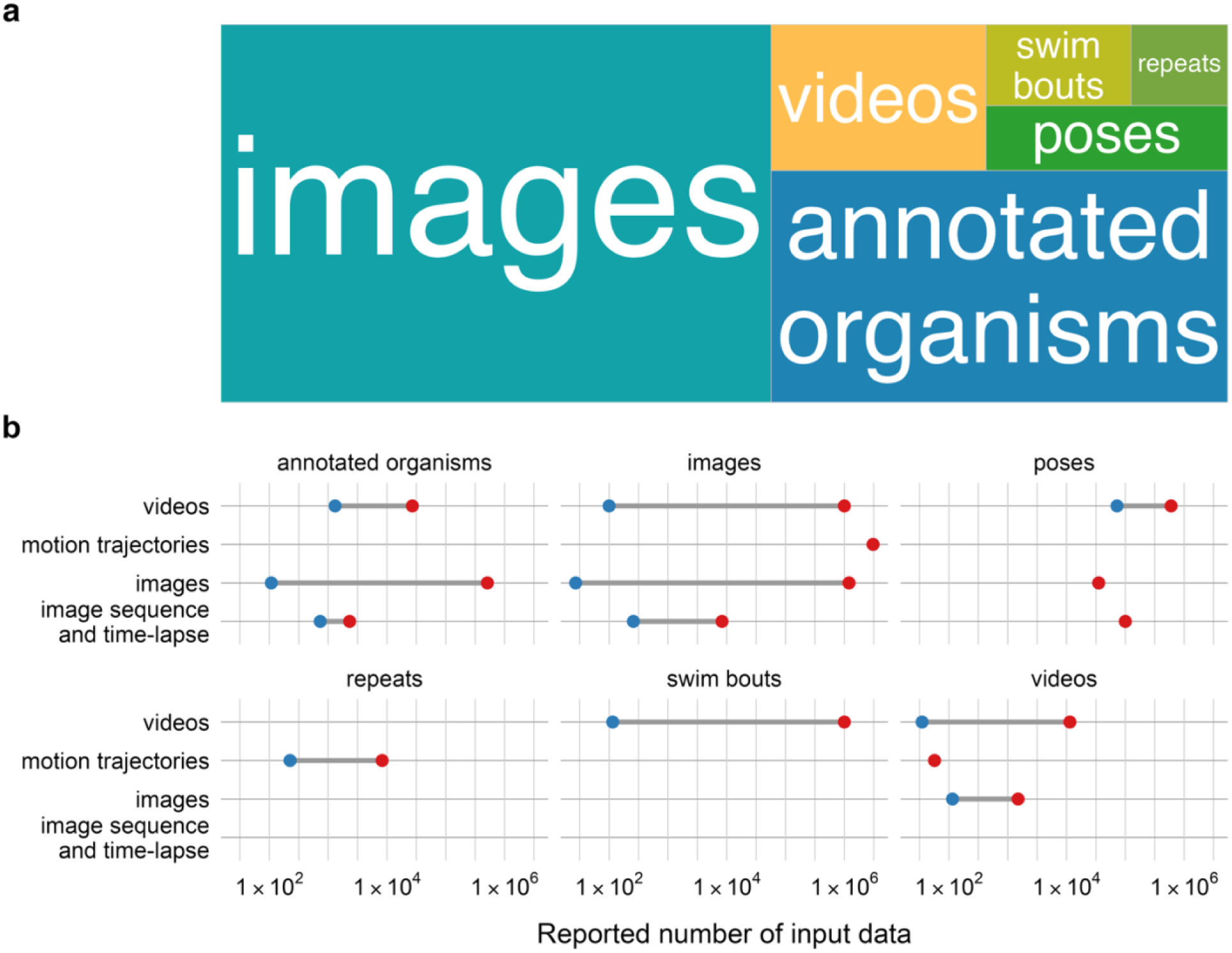
Overview of input data reported in analyzed studies. (a) Type of input data reported, size of boxes relative to the total number of publications reporting stated type, (b) Total number of input data used for model training according to the type of input data. The blue and red dots mark min. and max. number of input data on the range scale.

### Methods for behavior characterization and quantification

The portion of literature focused on analyzing behavior of model organisms*^29,31–34,36,37,39–42,44–46,49,51,52,54,55,57–59,62,71–73,79,81–86,88,91,92,97,102,104,107,108,114,126^* (43 papers, Table S3), rather than their detection, employed several main strategies for extracting behavior characteristics. The most common strategies included:

- Training algorithm to recognize defined model organisms’ shapes*^54,88,104,114^*, feeding behaviors*^31^*, body parts*^92,114^* or social interactions*^46^*,
- Tracking animals’ movement trajectories, shapes or poses to calculate kinematic movement parameters such as distance traveled, speed or stop duration *^33,40,51,71,72,85,97,107^*.
- Extract poses of animals for tracking experiments or to characterize behavior*^37,42,55,59,79,81,82,86^*,
- Extracting animal poses to classify or cluster behavior into groups (treated /naïve, wild-type/mutant, long-lived/short-lived) *^39,85,92,102,108^*,
- Tracking animals’ movement trajectories to cluster or classify movement patterns*^29,34,45,83,84^*.

These strategies either formulated the main core of the study characterizing organism’s behavior, discovered new movement patterns or were further used to assess behavior of model organisms under different physical and chemical conditions, inferring on animal lifespan or genotype. The behavioral characteristics investigated in analyzed papers are illustrated on Fig 10.

**Figure 10.**
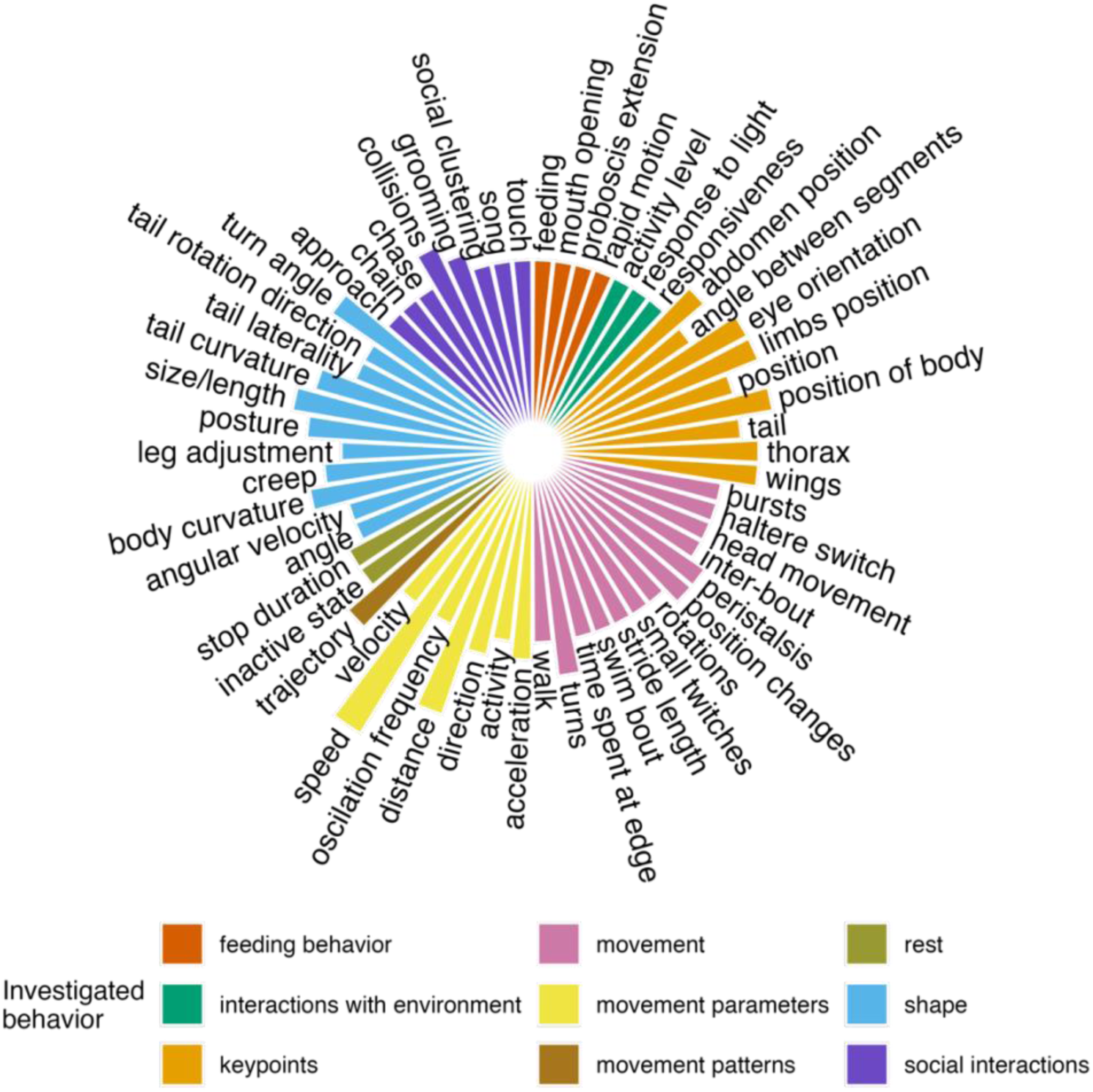
Overview of reported behavioral characteristics investigated during the analysis. The behavioral characteristics are color-grouped, and the size of bars represents relative frequency of studying behavioral characteristics.

The most widely used behavioral characteristics fall into the group of movement, movement parameters and organisms’ shapes (Fig 10). Features such as total distance, direction of travel or speed were readily calculated from tracking or pose estimation algorithms*^36,51,62,71,72,83,108^*. On the other hand, movement features and shapes were most commonly identified by the supervised learning algorithms, which learnt from manually labeled data*^31,46,52,54,57,85,92,102,107,114^*. Another group of behavioral characteristics widely used were keypoints*^37,41,42,55,59,72,81–84,97^*, that found applications in pose estimation models (Fig 10). Depending on the organism investigated, different sets of features are most informative. For instance, worms’ behavior was most commonly quantified in the form of shapes (such as coils, curvature, angle, length)*^32,33,36,45,57,59,72,91,92,104^*, whereas social behavior was almost exclusively investigated in flies and flying insects*^31,46^*. Feeding strikes and swimming patterns were the most frequent in studies on larval form of zebrafish*^34,40,108^*. The overwhelming majority of behavioral analysis was investigated on *D.* rerio*^29,34,40,49,82,83,85,107,108^*, D. melanogaster*^37,41,42,44,46,51,55,58,62,71–73,79,81,84,86,88,102^* and C. elegans*^32,33,36,45,57,59,72,91,92,104^* (84%).

### Preprocessing steps

Preprocessing techniques can be defined as steps taken to prepare raw data into correct format for analysis pipeline or model training. There is a wide range of techniques that are applied depending on core processing. In case of model training, for visual data analysis these include data labelling (for supervised ML and DL), converting data into tensors, splitting the data into train-val-test datasets, and many more, as shown in Table 4. The first 3 preprocessing steps have been omitted from the analysis below, as these form an obligatory core of preprocessing that computer scientists need to follow to develop AI-based model. In the following section, analysis will focus on the rest of preprocessing techniques, as grouped in Table 4.

**Table 4.**
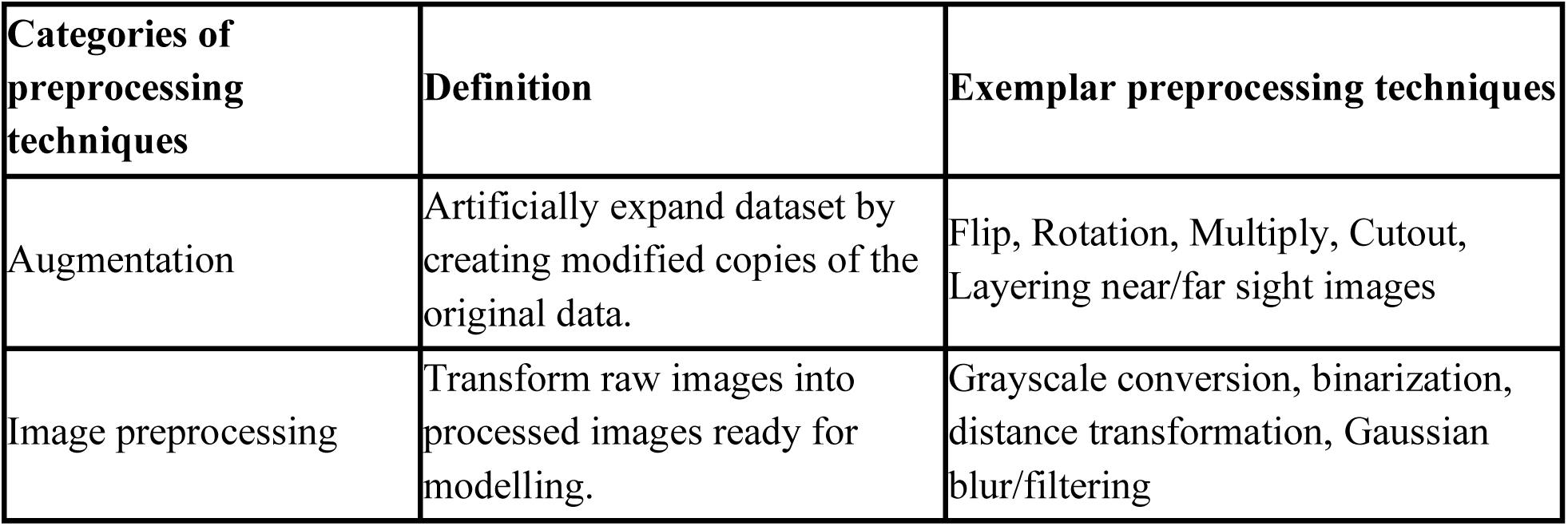
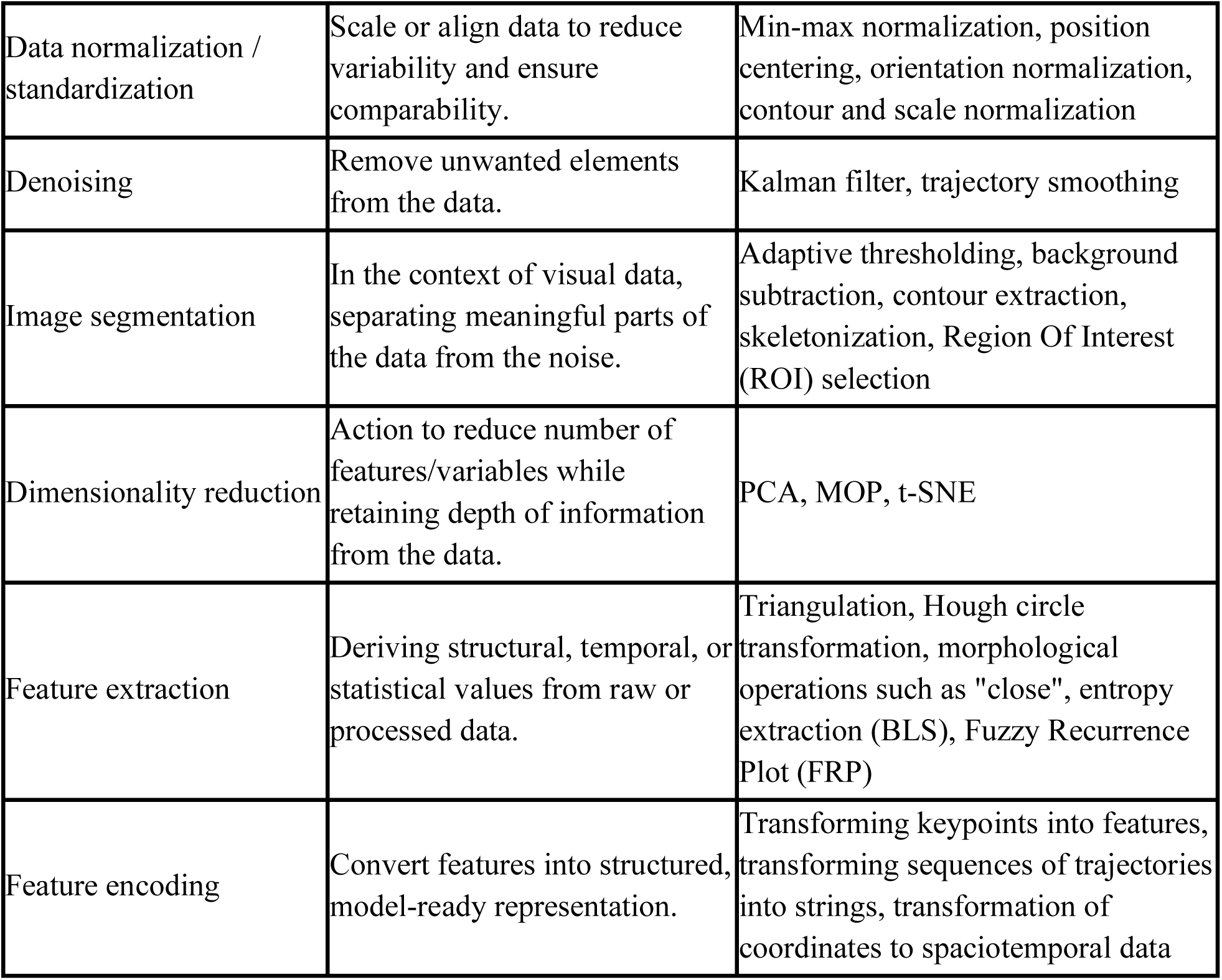
Categories of preprocessing techniques, exemplar techniques utilized in analyzed literature.

In the analyzed literature, there was a high degree of heterogeneity in reporting preprocessing steps. Some papers dedicated the whole subsection to describing steps undertaken to prepare the data for model training, sometimes accompanied with informative diagram, others barely mentioned said techniques. Hence, the data were summarized as a heatmap (Fig 11a) highlighting relative frequencies of preprocessing techniques undertaken according to the model type.

**Figure 11.**
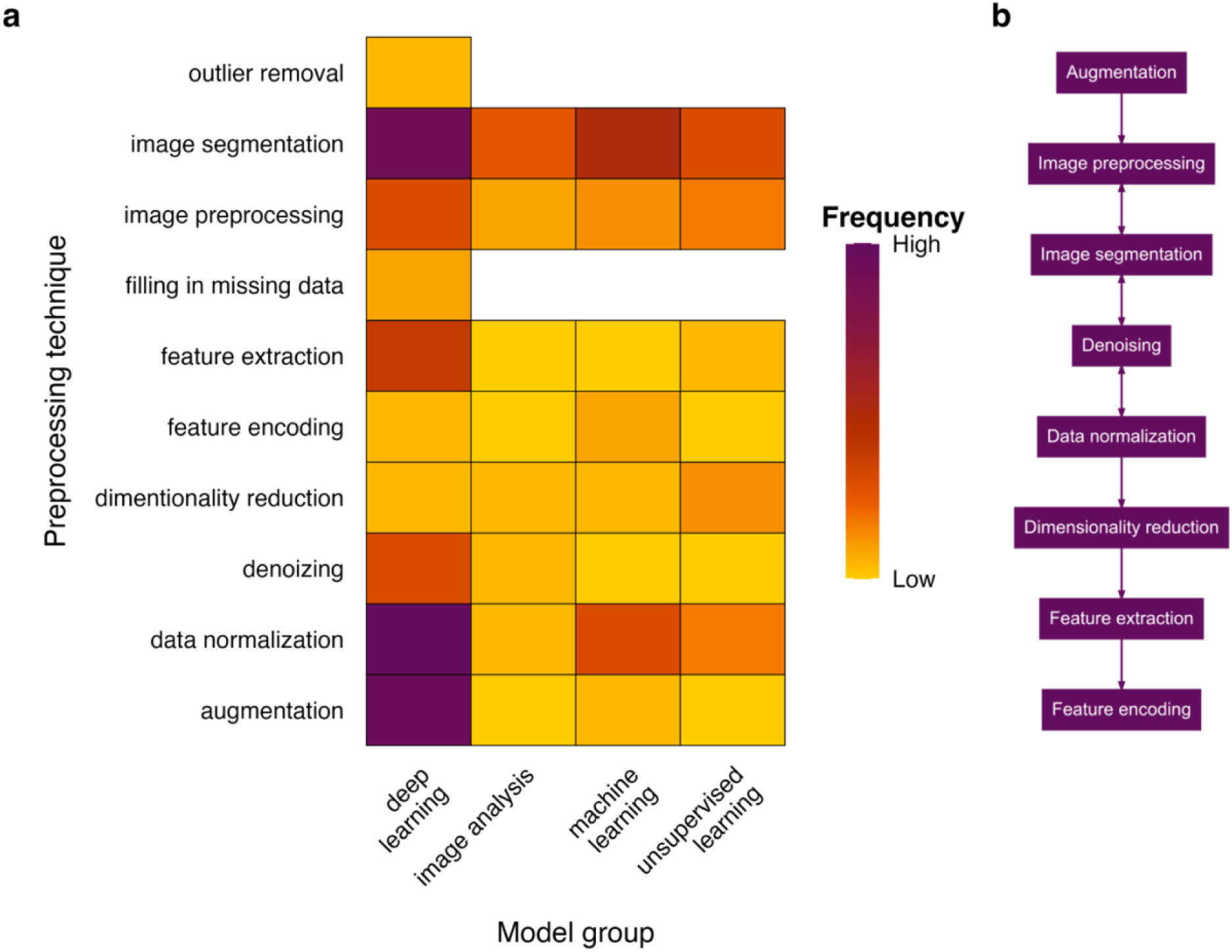
Reported preprocessing techniques according to different model types. Heatmap showing relative proportion of preprocessing techniques reported for each model group (a) and exemplary flow diagram of preprocessing steps (b).

The preliminary step in preprocessing was data augmentation*^30,42,47,53–56,66,67,69,75,76,80,92,94,101,110,111,116,120,123^*, most frequently used for DL models (Fig 11). Increasing the dataset size, by images resizing, rotating or reducing image clarity is a common practice in model training, especially for DL models, which require substantially more input data to achieve high performance. The most commonly used techniques within the image preprocessing category included Gaussian filtering*^59,60,79,118,124^* and grayscale conversion*^64,67,78^*. The former technique blurred the image to prevent overfitting at the same time putting emphasis on objects of interest. The latter technique allowed to reduce computational cost of image processing. Conversion of images from three-color scale such as RGB format (commonly used in camera recording settings) to single color scale such as greyscale preserves important information while reducing the memory space as well as processing time in model training. Next, many pipelines included image segmentation techniques, such as adaptive thresholding*^29,59,62–64^*, skeletonization*^36,45,50,105,106,125^* and background subtraction*^29,40,45,49,62,65,73,106,124,127^* (Fig 11). These techniques also serve the purpose of reducing data depth while preserving important features. Reducing computational cost and unnecessary depth of information optimizes model training time. The denoising and data normalization approaches were also reported in analyzed literature. Normalization included object centering*^58,121^* and scale normalization*^31,32,34,49,53,58,74,76,84,88,92,95,96^* whereas denoising techniques included trajectory smoothing*^82^*. Dimensionality reduction techniques were rarely reported at preprocessing level*^29,39,84,88^*. However, many unsupervised learning and multi-model pipelines included dimensionality reduction techniques at latter stages of the analysis*^45,49,74,79,84,85^*. Finally, feature extraction and feature encoding *^33,34,36,91,102,111,114^* were least commonly reported at this stage (Table 4, Fig 11).

Taken together, reporting of preprocessing steps was not standardized in analyzed literature. Possibly, due to the rapid advancements in AI techniques across a wide range of scientific fields, there is no standardization of methodology reporting. Moreover, many outlined pipelines utilized end-to-end software for tracking pose estimation (e.g. DeepSORT*^109^* and DeepLabCut*^55,84,97^*), that already included standard preprocessing steps as part of the model application. Interestingly, over 30% of pipelines utilizing DL models reported no preprocessing*^38,48,68,72,98,103,104,108^* or only data augmentation*^30,42,47,52,56,71,75,80,110,120,123^* (Table S4). Nonetheless, analyzed literature shows a clear preprocessing pipeline as shown in Figure 11b. Data is commonly augmented, images are then resized and color-scale converted followed by denoising and normalization techniques.

### Model architectures

Within a subgroup of DL models, the vast majority developed CNN models (85%) *^30,37,38,41,42,45–48,54–56,58–61,66–70,75,78–87,92–97,99,101,103–106,108–117,120–124,126,127^*, followed by Transformers (6%)*^72,73,98,119^*, RNN (4%), hybrid models of Transformer and CNN architecture elements (3%)*^57,118^* and autoencoders (3%)*^53,107^*. These groups have distinct architecture elements highlighted in Fig 12. The characteristics of CNN architectures include convolutional layers for spatial feature extraction and pooling operations for dimensionality reduction*^128^* (Fig 12a). They are effective for image analysis, especially for static frame-by-frame extractions. On the other hand, RNNs, especially LSTM models, are built from independent blocks that have input and output, with a gating mechanism that discards useless information over time*^129,130^* (Fig 12b). They excel in analyzing temporal information such as motion sequences*^129,130^*. Autoencoders are feedforward architectures that compress input to later decompress and reconstruct information*^131^* (Fig 12c). They are useful in unsupervised tasks to learn to detect behavioral patterns*^131^*. Finally, Transformers, being highly diverse in their architecture, can be unified to the following core architecture elements: self-attention module (weights importance of different elements within an input), positional encoding (inputs information on order or special arrangement of an input data) and task-specific head (maps transformed representation into the final output such as classification, detection etc.)*^132^* (Fig 12d).

**Figure 12.**
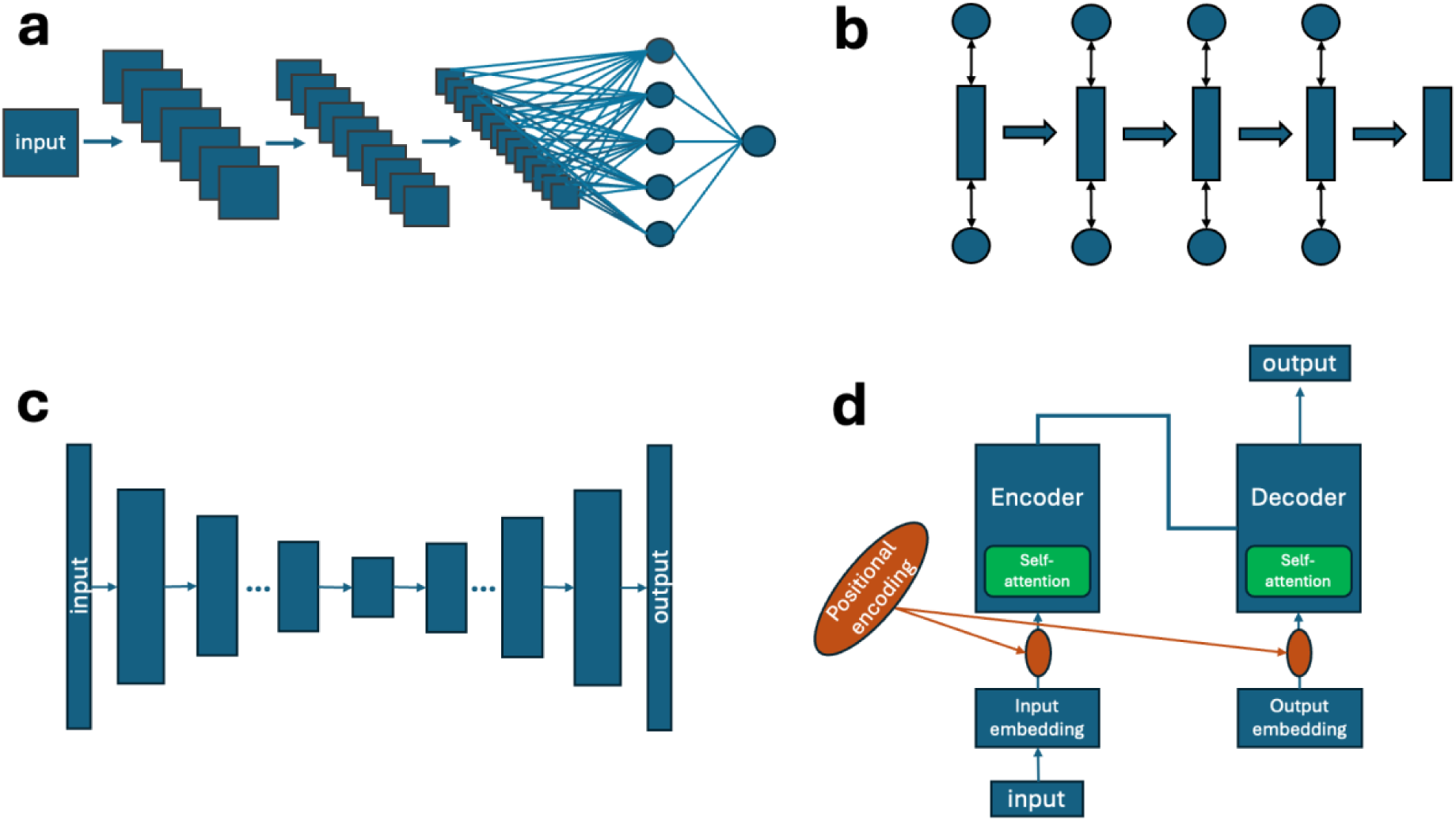
Graphical representation of basic architecture elements of four discussed architecture types: CNN (convolutional neural network) (a), RNN (recurrent neural network) (b), Autoencoder (c), Transformer (d).

Within analyzed literature, the most commonly reported architecture element was backbone*^37,41,42,46,48,55–57,66,70,75,77,79–81,87,92–94,96,98,101,104,108,110,111,114,116–118,120–124,126^* (Fig 13). Commonly utilized backbones include ResNet50*^37,42,55,66,77,94,104,108,114,126^*, CSPDarknet53*^70,87,101,111,117,120,122,123^* (used in YOLO models), and U-Net*^79,81,92,116^* (Table S5). In almost 50% of analyzed literature authors reported detailed model training such as type of optimizer used*^41,42,47,48,55,57–59,61,66,67,69,73,75,77,79,81,82,87,94,95,98–100,107,110,113–115,118,125,126^*, training epoch *^30,41,42,46–48,54,56–59,66,67,69,73,75,81,82,86,87,92–96,98–100,105,107,111,113–115,126^* and learning rate *^38,41,46–48,54,56–59,67,69,73,75,77,78,81,82,86,87,94,95,98–101,107,110,113,123,125^* (Table S5). The majority of models used Adam optimizer *^42,47,48,55,57–59,61,66,67,73,79,81,82,98,100,110,113–115,125,126^* (70%), 8 papers used Stochastic Gradient Descent (SGD) (24%)*^61,75,77,87,94,95,99,118^* and 2 papers used Root Mean Square Propagation (RMSProp) (6%)*^41,69^*. Training epoch ranged from 10 epoch (KAN Transformer *^73^*) to up to 125,000 epoch (Deep autoencoder *^107^*). Similarly, the learning rate of algorithms varied greatly, from 0.1*^123^* to 0.00001*^46,99^*. Furthermore, size of input data*^30,41,46,48,54,57–59,66,69,70,73,75,78,86,91–93,95,101,106,111,113,125,126^* was more frequently reported than the size of output data *^30,41,46,48,54,57–59,66,69,70,73,75,78,86,91–93,95,101,106,111,113,125,126^* (38% vs 19%, Fig 13). Memory footprint was reported only in 39% of papers*^37,40–42,46,47,58,73,75,77,79–83,93,95,98,105,108,110,111,117,118,120,122–124^* (out of which in all but 1*^111^* as GPU). Model neck was reported only in 16% of papers *^66,77,104,127^*, most commonly as Feature Pyramid Network (FPN) in Mask R-CNN *^66,77,104,127^* or other R-CNN derivatives*^60,96^*.

**Figure 13.**
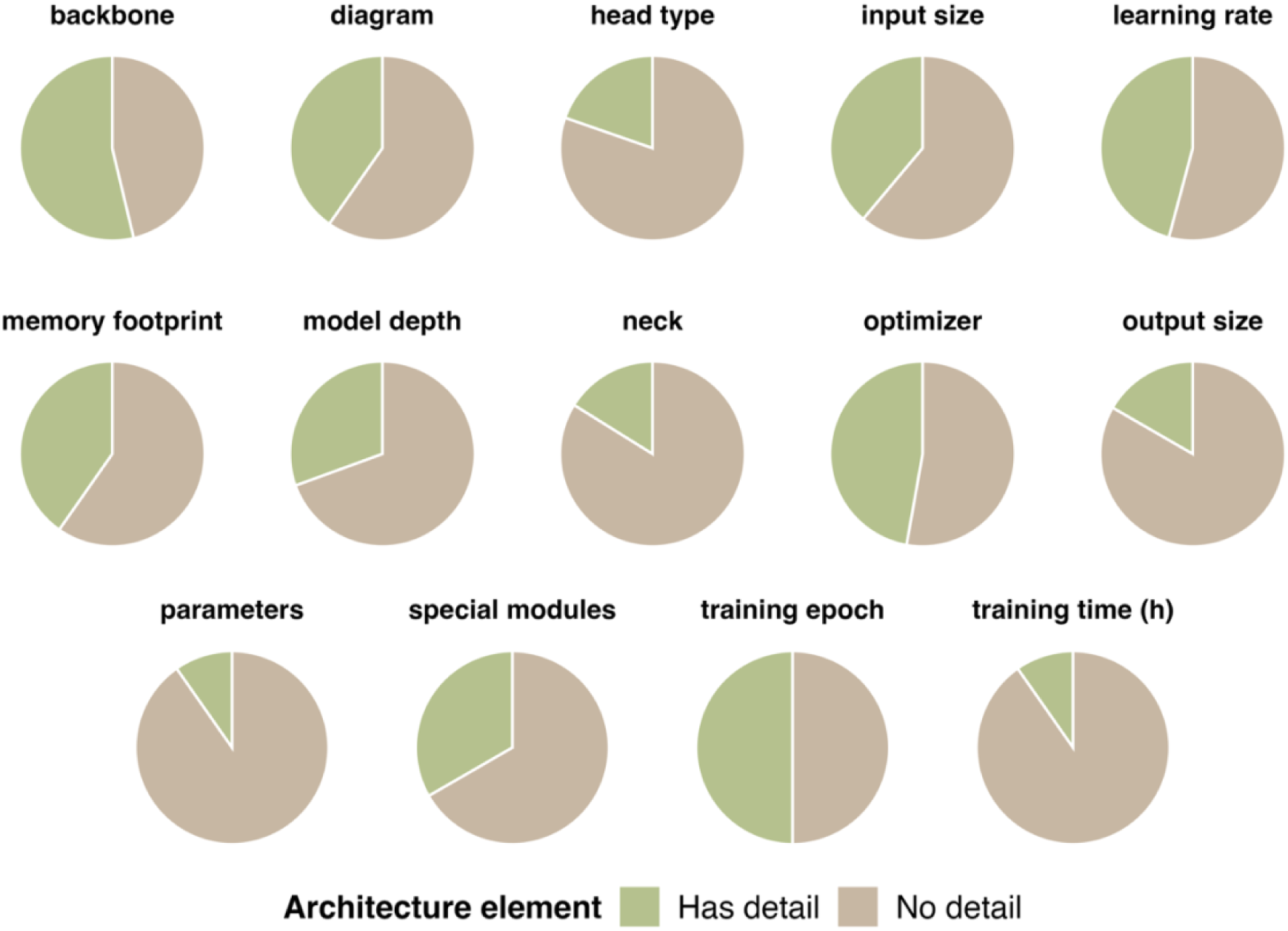
Visual representation of overall reporting of details on model architecture across analyzed literature.

Finally, the least reported model architecture details was the training time*^37,41,54,58,75,82,83^*, reported only in 10% of published research. As seen in Figure 13, the extent of architecture elements reported varies greatly across papers, highlighting an urgent need for standardization of AI-models in research. The raw table with all extracted information of models’ architecture can be reviewed in Table S5.

### Evaluation metrics

The key part of developing an algorithm is evaluating its performance. Without this step, the quality of the model can hardly be assessed. Within the analyzed literature, 46% of evaluated algorithms reported only 1 metric*^29,32,33,36–42,47–49,53–55,57–66,74,77–79,81–87,96,102,103,108,119,120,124,125^*. 19% and 17% of models reported 2*^35,43,44,46,69,75,79,93,97,101,104,105,108–110,125^* and 3*^52,67,70,72,76,80,88,91,99,111,113,115,118,122,123,127^* evaluation metrics, respectively, and the remaining 18% reported 4 or more metrics*^31,56,68,73,92,94,95,98,100,106,107,112,114,116,117,121,126^* (Table S6).

Figure 14 shows the frequency of reported evaluation metrics across different model groups: DL, ML, unsupervised learning, and others. For clarity of visualization, only metrics reported more than once across all research papers were shown on the figure. Accuracy*^31,36,38,45,53,54,56,57,60,62,64,65,67–69,74,76,78,88,92,93,97,99–102,104,108,112–117,121,124,126^* was the most commonly reported evaluation metric in DL, ML and other models (Fig 14). Unsupervised learning models, due to their distinct application and methodology, reported quality of their models visually, on graphs showing distinct clusters of data*^39,49^*, statistically*^29,33^* or stating how much variability was explained by the model*^32,79^*. DL models show greatest variability in reported evaluation metrics, both because they were used for a wide group of tasks, and because the total number of DL models is significantly greater than other groups, introducing variability across reported research. While accuracy, precision, and recall, among others, are common evaluation metrics for detection and classification tasks *^31,36,38,45,53,54,56,57,60,62,64,65,67–69,74,76,78,88,92,93,97,99–102,104,108,112–117,121,124,126^*, for pose estimation and tracking metrics such as error*^42,44,58,82,83^*, MAE*^37,73,108^*, MOTA*^73,109,125^* and MOTP *^73,109,125^* are preferred. Unfortunately, the heterogeneity of reported evaluation metrics (Table S6) is mirrored by a range of values reported. The ranges of the three most commonly reported metrics for supervised learning and deep learning models are shown in Table 5.

**Figure 14.**
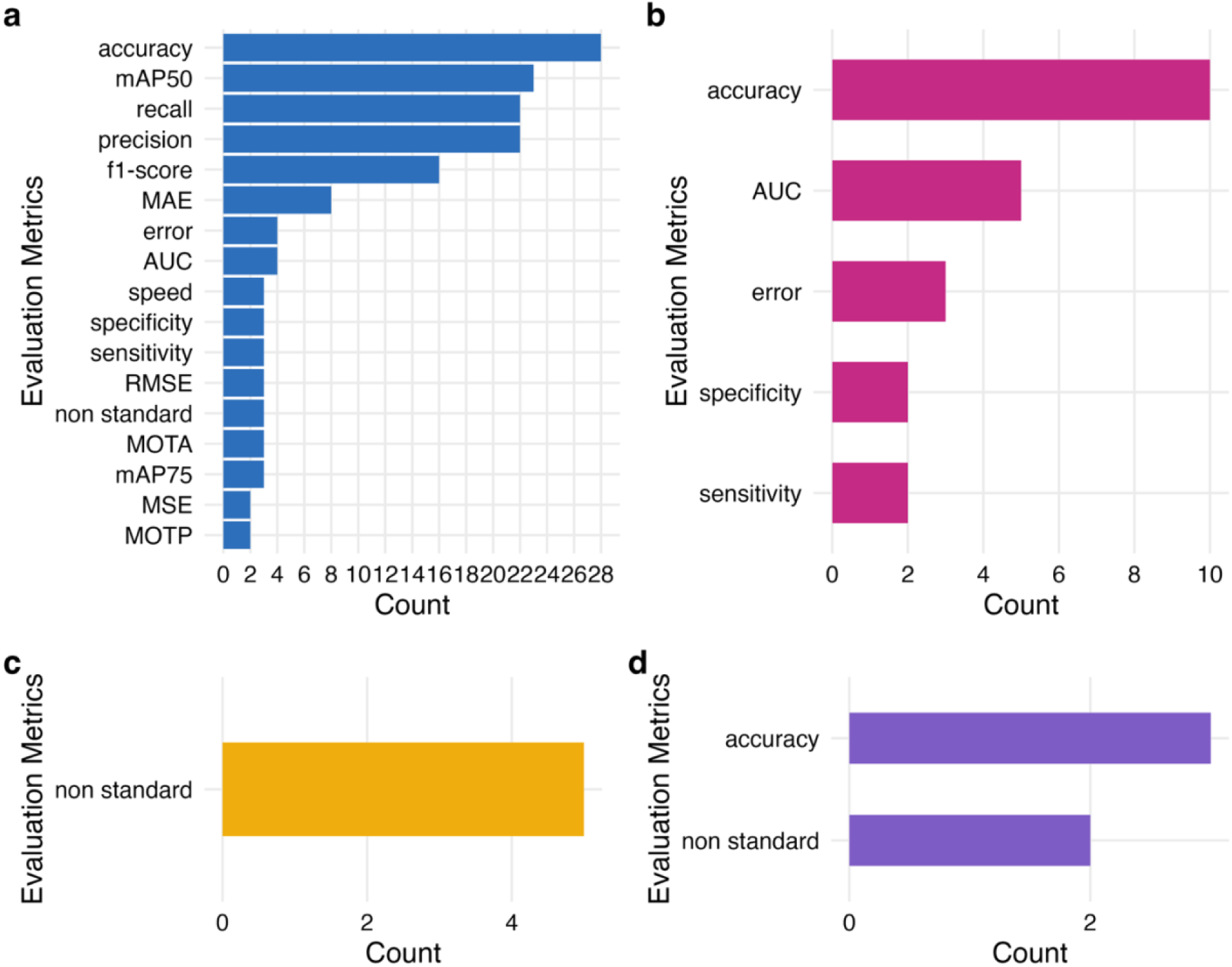
Overview of evaluation metrics reported for four groups of algorithms: Deep Learning (a), Supervised Learning (b), Unsupervised Learning (c) and Other (d) (incl. image analysis and statistical models). mAP50 – mean average precision at an intersection over union threshold of 0.5, MAE – mean average error, AUC – area under the curve, RMSE – root mean square error, MOTA – multi-object tracking accuracy, mAP75 - mean average precision at an intersection over union threshold of 0.75, MSE – mean standard error, MOTP – multi-object tracking precision.

**Table 5.**
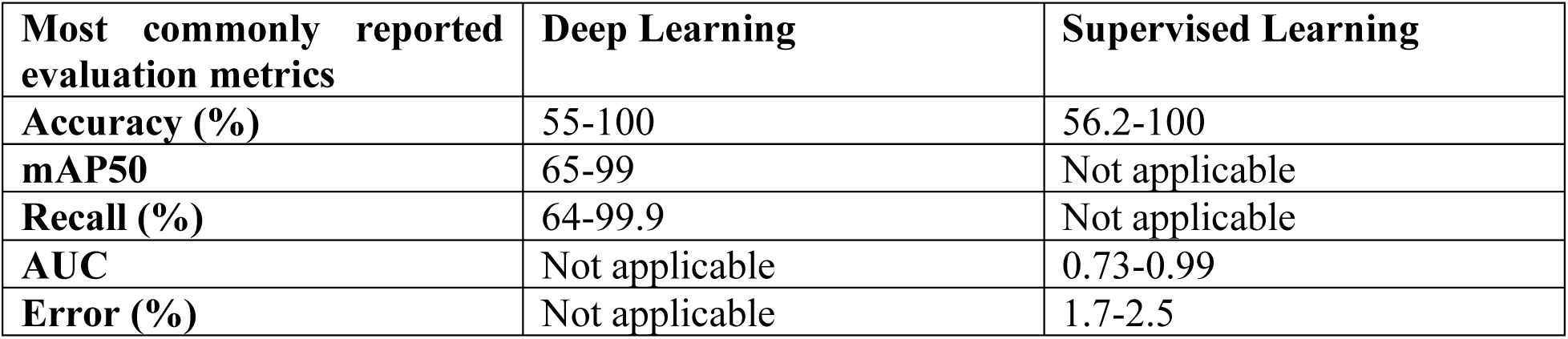
Metric values for the three most commonly reported evaluation metrics in Deep Learning and Supervised Learning models. AUC – area under the curve.

## Discussion

Invertebrate model organisms such as *C. elegans* or *D. melanogaster* remain among the most widely used systems in biology and biomedical research*^133^*. Their physiology, behavior, and genetics are well established through decades of investigation*^133,134^*, providing a robust foundation for experimental studies. These organisms, along with other invertebrates and an increasing number of larval models, offer a distinct advantage over vertebrates such as mice or rats in the context of high-throughput screenings and early-stage research*^134^*. Behavioral analysis is an emerging, powerful approach in those types of studies, enabling the systematic assessment of functional outcomes that corelate organism-level phenotypes with molecular and cellular mechanisms, and phenomena *^20^*. With the rapid adoption of AI-based methods by the scientific community, the potential of behavioral analysis in these model organisms has become a subject of active investigation.

The growing interest is reflected in a steady rise in publications over the last 15 years (Fig 2), spanning multiple biological disciplines. Fields such as neuroscience, epidemiology, and agriculture (Fig 6) started to adopt AI-based methods to study behavior, using over 15 different model organisms (Fig 5). In the last decades authors from over 34 countries shared their research in conference proceedings or peer-reviewed journals (Fig 3, Table S2), out of which 74% was published by journals with impact factor of 3 and more (Fig 4). This global trend, in terms of geography and research fields, further highlights that the field of AI-based methods in behavioral analysis is of ongoing investigation and current relevance, posing promising direction of future research.

Characterizing behavior is the first step in analysis, as it determines algorithm’s tasks and methodologies applied. Model tasks can be broadly categorized into three groups. First, detection and classification algorithms categorize behaviors based on predefined criteria such as locomotion parameters (e.g. distance traveled, speed) or morphological features (e.g. body curvature, coiling). These approaches have traditionally relied on classical machine learning models such as RF, SVM, or KNN, valued for their relatively low computational demands and interpretability*^135^*. However, they depend on researcher-defined categories, which may overlook ambiguous or subtle behaviors. DL models, such as YOLO or EfficientNet, have also found applications in these tasks providing greater capacity for analysis of images as input data, at the expense of requirement for larger computational power and training data resources*^136^*. A second category are pose estimation and tracking algorithms, which extract keypoints (e.g. head, limb or body coordinates) to describe movement kinematics. Such models are inherently objective, capturing raw motion data without imposing predefined behavioral labels, and have been used to quantify fine-scale phenotypes such as *D. melanogaster* body kinematics *^55^* or behavioral changes of nematodes exposed to essential oils *^97^*. Finally, unsupervised clustering and dimensionality reduction algorithms enable the discovery of novel behavioral motifs, such as swimming patterns *^34^* or prey-capture swim-bouts *^49^* in *D. rerio* larvae. These data-driven approaches minimize bias from predefined labels and are highly exploratory, but they pose significant challenges for biological interpretation and require careful validation.

Taken together, each class of methods offers distinct advantages and limitations, ranging from the subjectivity of manual annotation to the interpretability challenges of unsupervised learning*^137^*. These trade-offs are summarized in Figure 15, which highlights the specific strengths and weaknesses of individual techniques. For example, historically used manual observation does not require specialized equipment and leads to biologically relevant interpretation of behavior, however, it suffers from low efficiency, throughput and observer bias. Semi-automatic techniques improve scalability but reduce behavior to simple parameters. Finally, AI-based techniques aim to eliminate mentioned limitations, providing various approaches to behavioral analysis. Classical ML has proven particularly useful in analyses of smaller datasets, especially when predicting binary outcomes*^135^*. However, their main limitation lies in their low performance when it comes to recognizing complex and nonlinear behavioral patterns. Compared to classical ML, DL can process highly dimensional, unstructured data and yield more detailed interpretations*^137^*. However, due to the complexity of their architecture, models are less explainable, as developers do not directly program how models learn and hence do not know how models arrived at their conclusions*^136^*. This problem is referred to as “the black-box”. Finally, unsupervised learning models offer a method to discover novel patterns, and study behavior without prior behavioral assumptions. This comes at the expense of model complexity and previously mentioned challenges with placing and evaluating the findings in the biological context*^137^*.

**Figure 15.**
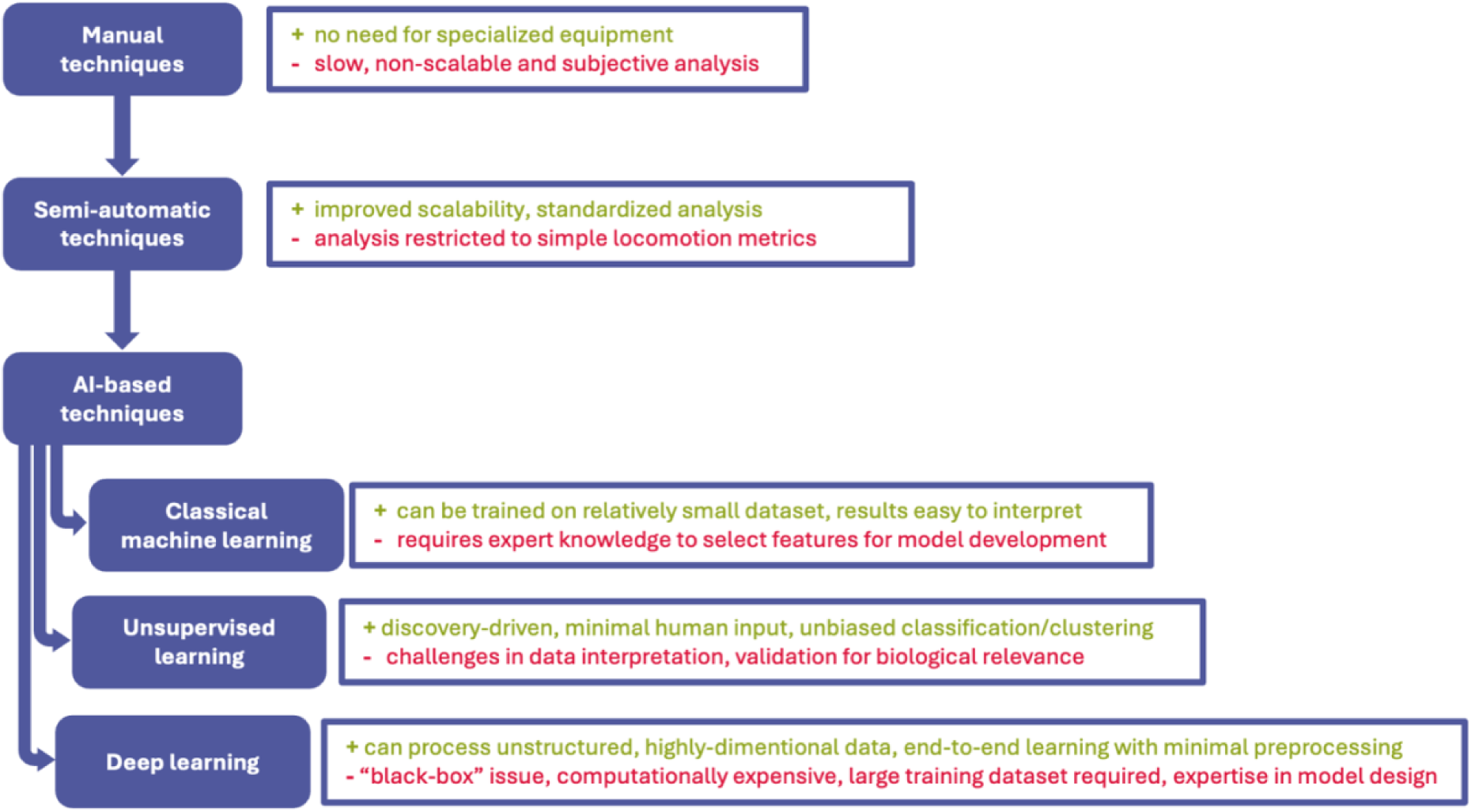
Advantages and disadvantages of methods used in behavioral analysis.

The analysis of published literature indicates that since 2021, DL models started to dominate the landscape of behavioral analysis in invertebrates and larval model organisms (Fig 7a). This trend can be attributed to several converging factors. First, the increasing availability of GPU computing has enabled the scaling of DL architectures, allowing them to be trained and applied to large and complex datasets. Second, the emergence of user-friendly frameworks such as YOLO*^138^* has lowered the technical barrier for adoption, providing researchers without extensive computational expertise with accessible tools for detection, classification, and tracking. For example, YOLO hosted on Roboflow offers an intuitive online interface where users can upload and annotate images and train models with minimal coding requirements*^139^*. Finally, the development of more advanced, open-source platforms such as DeepLabCut*^37^* and SLEAP*^81^* has provided flexible tools for complex pose estimation and behavioral analyses. The impact of such platforms is evident in the widespread use of DeepLabCut, whose original paper, published in 2020 *^37^*, has been cited more than 4,300 times (Nature, sampled on 14/10/2025), underscoring its role in enabling reproducible research across diverse model organisms and experimental conditions.

Within the group of DL models there has been an emergence of several subgroups worth mentioning due to their potential in behavioral research. Most dominant DL-based group, CNN models, are characterized by high efficiency and automation of the analysis process, especially in the context of visual data processing*^128^*. However, their limitation is their inability to process temporal dependencies, which limits them in motion sequence prediction tasks*^128^*. In response to this limitation, sequential architectures, including RNN and transformers, are increasingly being used to enable the analysis of dynamic changes over time. RNN models, despite their simple architecture, work well for trajectory prediction and motion micro-pattern detection*^129^*. Transformers, on the other hand, allow for the analysis of complex spatiotemporal contexts, making them particularly useful for tasks involving the recognition of intentional behaviors and motivational states*^132^*. Despite highlighted advantages of both RNN and Transformer architectures, they are only emerging in the field and comparative studies between different architectures across model tasks will reveal their advantages and disadvantages in behavioral analysis of invertebrate and larval model organisms.

Described DL architectures approach behavioral analysis either detecting and classifying patterns or poses or by tracking model organism’s movement. The majority of models fall under the category of supervised learning, whether classical or DL, meaning they require manually labeled images or videos for training *^137^*. This approach is time-consuming and relies on experts’ observation. Moreover, in most cases, the algorithm learns on a defined set of data, creating a loophole for biased analysis when investigating previously unencountered behavioral patterns. Pereira et al approached overcoming this limitation by developing a CNN model that identifies useful images for labeling and allows users to manually annotate those images via graphical user interface *^42^*. On the other hand, Gunel et al incorporated a module that iteratively reprojects 3D poses of animals to automatically detect and correct 2D errors, which in turn allows for further training the network without user intervention *^41^*. Both approaches show how models can constantly improve on precision, accuracy and other evaluation metrics, aiming to expand their capabilities for an increasingly wider set of behavioral patterns*^42^*. Implementing active learning modules in models investigating subjects as dynamic and complex as behavior would improve its applicability across model organisms and experimental designs, making them more adaptable and reliable tools in scientific research.

An alternative approach to overcome issues resulting from training models on predefined labels is the use of unsupervised learning. The method, by definition, does not require manual labels and learns patterns in data without human supervision*^137^*. Analysis is therefore not influenced by scientists’ understanding of behavior, allowing for discoveries of novel or even reconstructing known patterns. However, at the same time, model validation, to place findings in a biologically relevant context is essential. Some research papers proposed combining unsupervised approaches with supervised learning *^39^*or deep learning *^45^* to cluster behavioral patterns and later classify the behavior. Other developed complex pipelines involving tracking or pose estimation (via DL), combined with classification and clustering or dimensionality reduction techniques *^84,85,140^*. These models are rarely implemented, possibly due to their complexity and requirement for collaboration between wet-lab with deep behavioral expertise on model organism, with ML experts. Their potential at the interface of biology and computational science remains to be discovered.

Current approaches to modeling invertebrate behavior rely predominantly on supervised learning, both classical and deep learning, which are trained on manually labeled data. This dependence on human annotation introduces bias, as models learn from the subjective knowledge and assumptions of researchers, limiting their ability to provide fully objective analyses. Pose estimation has become a widely used framework for quantifying movement and offers an objective representation of behavior that can be classified, clustered or interpreted otherwise. However, datasets remain highly imbalanced, with frequent locomotor patterns overrepresented while rare but biologically important behaviors can be undetected or entirely missed. Developing behavioral ontologies and open, standardized datasets could help overcome this issue by enabling the integration of learned behavioral patterns across studies, similar to the role of ImageNet or COCO in computer vision. Reporting datasets, as Li et al *^72^* and Pereira et al *^81^*, is the first step towards transparency and forms foundation for future datasets integration to group behavior and classify behavioral patterns for different model organisms.

Another major limitation lies in the heterogeneity of reported data. The studies differ in the form of reporting number of input samples, recording setups (incl. frame rate, resolution, and lighting), preprocessing pipelines, documentation of model architectures and training procedures. These inconsistencies make it difficult to reproduce findings and to directly compare models. The limited number of comparative studies that benchmark different approaches is likely a direct consequence of this fragmentation. To address these challenges, we propose the introduction of a standardized reporting template for behavioral modeling studies (File S1). Such framework should include detailed documentation of experimental design, input data, preprocessing steps, and model training details, thereby improving transparency and reproducibility, and ultimately facilitating cross-laboratory comparison and cumulative progress in the field. The proposed workflow for reporting these elements can be reviewed in Figure 16, and complete reporting data sheet can be viewed in File S1.

**Figure 16.**
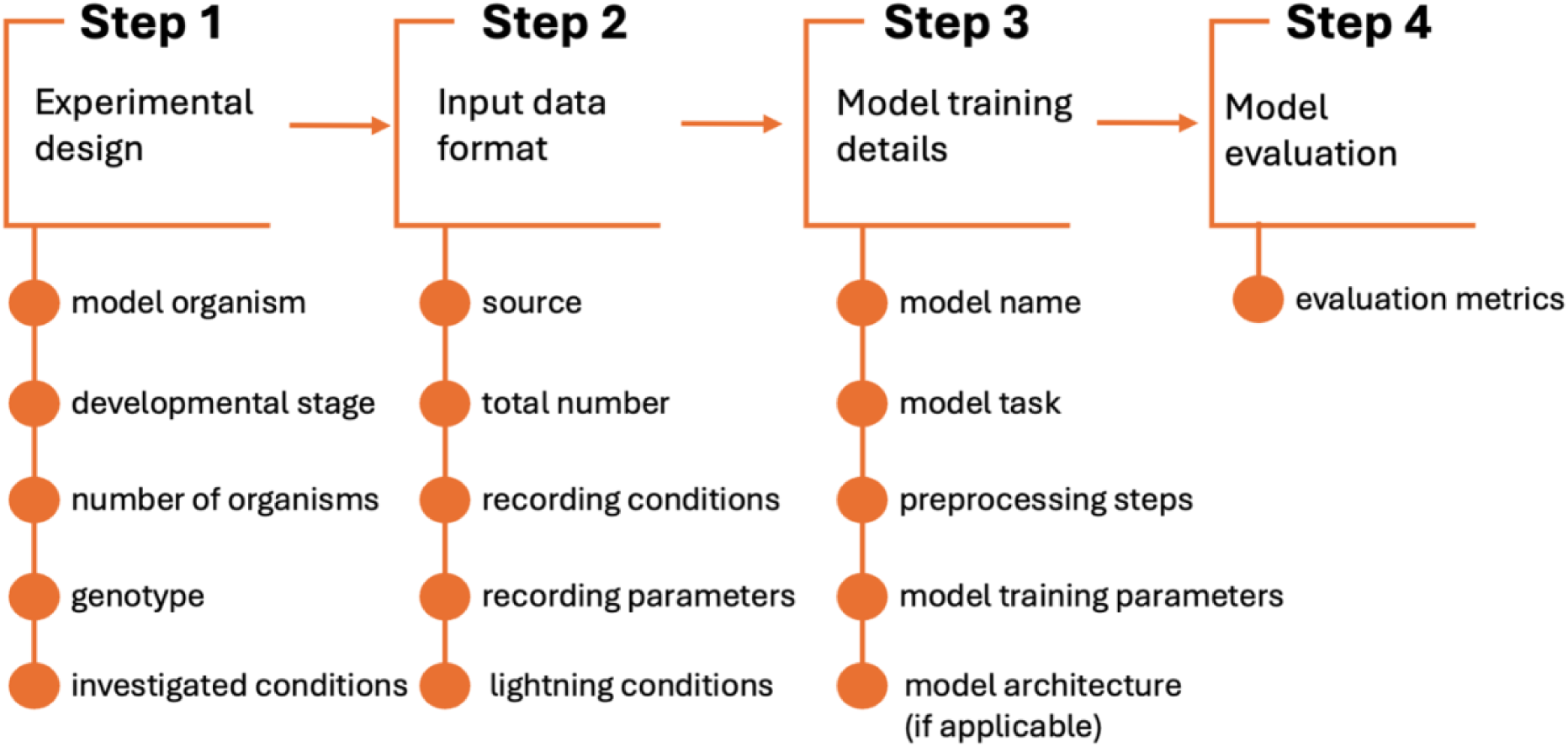
Proposed workflow for recording key information on model design and evaluation for behavioral studies on invertebrate and larval model organisms.

Looking ahead, behavior is increasingly recognized as a valuable tool across scientific disciplines, from deconstructing animal actions to conducting high-throughput toxicological screens and early-stage drug discovery. This complex and multidimensional aspect of physiology offers substantial potential for quantitative analysis. Advances in computational science and AI-based methods have enabled diverse models for behavioral analysis in invertebrate and larval organisms. These models detect poses, track movement, and classify behaviors, providing powerful high-throughput tools. Each approach has unique strengths and limitations, as outlined in Figure 15, and AI-driven behavioral analysis in invertebrates remains underexplored.

The emergence of Transformer, RNN, and hybrid architectures underscores the rapid evolution of computational approaches. Integrating multimodal data presents a promising direction for enhancing interpretability and analytical depth. To ensure reliability and biological relevance, future models must be explainable, transparent, and adaptable to experimental design. Standardized behavioral data reporting will be essential to enable collaboration and cross-disciplinary progress. Continued development of AI-based behavioral analysis holds significant potential to advance toxicology, pharmacology, and fundamental research, driving a deeper understanding of animal behavior across model systems.

## Data availability

All data collected for the systematic review are included in this published article and supplementary materials.

## Author Contributions

Conceptualization, N.P.; Methodology N.P. and Z.S; Literature review, Z.S. and N.P.; Data analysis and visualizations, Z.S.; Manuscript writing, Z.S., N.P., A.J., M.B., T.M.; Manuscript review and editing, A.J, M.B., T.M.; Supervision, A.J.; Project administration, A.J.; Funding acquisition, A.J. All authors have read and agreed to the published version of the manuscript.

## References

1. Obafemi, O. T. et al. Animal models in biomedical research: Relevance of Drosophila melanogaster. Heliyon 11, (2025).

2. Pastorino, P., Prearo, M. & Barceló, D. Ethical principles and scientific advancements: In vitro, in silico, and non-vertebrate animal approaches for a green ecotoxicology. Green Analytical Chemistry 8, 100096 (2024).

3. Strähle, U. et al. Zebrafish embryos as an alternative to animal experiments-A commentary on the definition of the onset of protected life stages in animal welfare regulations. Reproductive Toxicology 33, 128–132 (2012).

4. Sison, M. & Gerlai, R. Behavioral performance altering effects of MK-801 in zebrafish (Danio rerio). Behavioural Brain Research 220, 331–337 (2011).

5. Smith, L. L. et al. Strengths and limitations of morphological and behavioral analyses in detecting dopaminergic deficiency in Caenorhabditis elegans. Neurotoxicology 74, 209–220 (2019).

6. Ménard, G., Rouillon, A., Cattoir, V. & Donnio, P. Y. Galleria mellonella as a Suitable Model of Bacterial Infection: Past, Present and Future. Front Cell Infect Microbiol 11, 782733 (2021).

7. Marena, G. D., Thomaz, L., Nosanchuk, J. D. & Taborda, C. P. Galleria mellonella as an Invertebrate Model for Studying Fungal Infections. Journal of Fungi 2025, Vol. 11, *Page 157* 11, 157 (2025).

8. Asai, M., Li, Y., Newton, S. M., Robertson, B. D. & Langford, P. R. Galleria mellonella-intracellular bacteria pathogen infection models: The ins and outs. FEMS Microbiol Rev 47, (2023).

9. Ratner, M. H. & Farb, D. H. Probing the Neural Circuitry Targets of Neurotoxicants In Vivo Through High Density Silicon Probe Brain Implants. Frontiers in Toxicology 4, (2022).

10. Rosa, J. G. S., Lima, C. & Lopes-Ferreira, M. Zebrafish Larvae Behavior Models as a Tool for Drug Screenings and Pre-Clinical Trials: A Review. Int J Mol Sci 23, (2022).

11. Maximino, C. et al. Measuring anxiety in zebrafish: A critical review. Behavioural Brain Research 214, 157–171 (2010).

12. Jarema, K. A. et al. Developmental Neurotoxicity and Behavioral Screening in Larval Zebrafish with a Comparison to Other Published Results. Toxics 10, 256 (2022).

13. Jiang, Y., Chen, J., Wu, Y., Wang, Q. & Li, H. Sublethal Toxicity Endpoints of Heavy Metals to the Nematode Caenorhabditis elegans. PLoS One 11, 148014 (2016).

14. Egnor, S. E. R. & Branson, K. Computational Analysis of Behavior. in ANNUAL REVIEW OF NEUROSCIENCE (ed. Hyman, S. E.) vol. 39 217–236 (2016).

15. Demir, E. & Dickson, B. J. fruitless splicing specifies male courtship behavior in Drosophila. Cell 121, 785–794 (2005).

16. Desland, F. A., Afzal, A., Warraich, Z. & Mocco, J. Manual versus Automated Rodent Behavioral Assessment: Comparing Efficacy and Ease of Bederson and Garcia Neurological Deficit Scores to an Open Field Video-Tracking System. J Cent Nerv Syst Dis 6, JCNSD.S13194 (2014).

17. Winter, M. J. et al. Validation of a larval zebrafish locomotor assay for assessing the seizure liability of early-stage development drugs. J Pharmacol Toxicol Methods 57, 176–187 (2008).

18. Luxem, K., et al. Open-Source Tools for Behavioral Video Analysis: Setup, Methods, and Development. https://arxiv.org/pdf/2204.02842 (2022).

19. Berman, G. J. Measuring behavior across scales. BMC Biol 16, 1–11 (2018).

20. Adjé, E. A., Ahouandjinou, A. S. R. M., Delmaire, G., Roussel, G. & Houndji, R. V. Advancements in Video-Based Insect Tracking: A Bibliometric Analysis to A Short Survey. in ACM International Conference Proceeding Series 75–82 (Association for Computing Machinery, 2023). doi:10.1145/3635118.3635130.

21. Egnor, S. E. R. & Branson, K. Computational Analysis of Behavior. Annual Reviews of Neuroscience 39, 217–236 (2016).

22. Robie, A., Seagraves, K., Egnor, S. & Branson, K. Machine vision methods for analyzing social interactions. Journal of Experimental Biology https://journals.biologists.com/jeb/article-abstract/220/1/25/33433 (2017).

23. Moulin, T. C., Covill, L. E., Itskov, P. M., Williams, M. J. & Schiöth, H. B. Rodent and fly models in behavioral neuroscience: An evaluation of methodological advances, comparative research, and future perspectives. Neurosci Biobehav Rev 120, 1–12 (2021).

24. Mathis MW & Mathis, A. Deep learning tools for the measurement of animal behavior in neuroscience. Curr Opin Neurobiol 60, 111 (2020).

25. Lima, M., Damascena de Almeida Leandro, M., Valero, C., Coronel, L. & Bazzo, C. Automatic detection and monitoring of insect pests—A review. Agriculture https://www.researchgate.net/profile/Maria-Elisa-Leandro/publication/341277176_Automatic_Detection_and_Monitoring_of_Insect_Pests-A_Review/links/5eb9b15d299bf1287f7fb004/Automatic-Detection-and-Monitoring-of-Insect-Pests-A-Review.pdf (2020).

26. Pereira, T., Shaevitz, J. & Murthy, M. Quantifying behavior to understand the brain. Nat Neurosci https://www.nature.com/articles/s41593-020-00734-z (2020).

27. Page, M. J. et al. The PRISMA 2020 statement: an updated guideline for reporting systematic reviews. BMJ 372, (2021).

28. Cooke, A., Smith, D. & Booth, A. Beyond PICO: The SPIDER tool for qualitative evidence synthesis. Qual Health Res 22, 1435–1443 (2012).

29. Girdhar, K., Gruebele, M. & Chemla, Y. R. The behavioral space of zebrafish locomotion and its neural network analog. PLoS One 10, (2015).

30. Stern, U., He, R. & Yang, C.-H. Analyzing animal behavior via classifying each video frame using convolutional neural networks. Sci Rep 5, (2015).

31. Shamur, E. et al. Automated detection of feeding strikes by larval fish using continuous high-speed digital video: A novel method to extract quantitative data from fast, sparse kinematic events. Journal of Experimental Biology 219, 1608–1617 (2016).

32. Huang, L., Kim, H., Furst, J. & Raicu, D. A Run-Length Encoding Approach for Path Analysis of C. elegans Search Behavior. Comput Math Methods Med 2016, (2016).

33. Perni, M. et al. Massively parallel C. elegans tracking provides multi-dimensional fingerprints for phenotypic discovery. J Neurosci Methods 306, 57–67 (2018).

34. Marques, J. C., Lackner, S., Félix, R. & Orger, M. B. Structure of the Zebrafish Locomotor Repertoire Revealed with Unsupervised Behavioral Clustering. Current Biology 28, 181–195.e5 (2018).

35. Huang, L. et al. Potential of visible and near-infrared hyperspectral imaging for detection of Diaphania pyloalis larvae and damage on mulberry leaves. Sensors 18, (2018).

36. Javer, A., Ripoll-Sánchez, L. & Brown, A. E. X. Powerful and interpretable behavioural features for quantitative phenotyping of Caenorhabditis elegans. Philosophical Transactions of the Royal Society B: Biological Sciences 373, (2018).

37. Mathis, A., Mamidanna, P., Cury, K. & … T. A. DeepLabCut: markerless pose estimation of user-defined body parts with deep learning. Nat Neurosci 21, 1281– 1289 (2018).

38. Aqil, M. et al. Training of convolutional neural network using transfer learning for Aedes Aegypti larvae. Telkomnika 16, 1894–1900 (2018).

39. Rudolf, J., Dondorp, D., Canon, L., Tieo, S. & Chatzigeorgiou, M. Automated behavioural analysis reveals the basic behavioural repertoire of the urochordate Ciona intestinalis. Sci Rep 9, (2019).

40. Štih, V., Petrucco, L., Kist, A. M. & Portugues, R. Stytra: An open-source, integrated system for stimulation, tracking and closed-loop behavioral experiments. PLoS Comput Biol 15, (2019).

41. Günel, S. et al. Deepfly3D, a deep learning-based approach for 3D limb and appendage tracking in tethered, adult Drosophila. Elife 8, (2019).

42. Pereira, T. D. et al. Fast animal pose estimation using deep neural networks. Nat Methods 16, 117–125 (2019).

43. Bornhorst, J., Nustede, E. J. & Fudickar, S. Mass Surveilance of C. elegans-Smartphone-Based DIY Microscope and Machine-Learning-Based Approach for Worm Detection. Sensors 19, (2019).

44. Wu, S. et al. Fully automated leg tracking of Drosophila neurodegeneration models reveals distinct conserved movement signatures. PLoS Biol 17, (2019).

45. Jeong, I. S., Lee, S. R., Song, I. & Kang, S. H. A Biological Monitoring Method based on the Response Behavior of Caenorhabditis Elegans to Chemicals in Water. Journal of Environmental Informatics 33, 47–55 (2019).

46. Jiang, Z., Chazot, P. L., Celebi, M. E., Crookes, D. & Jiang, R. Social Behavioral Phenotyping of Drosophila With a 2D–3D Hybrid CNN Framework. IEEE Access 7, 67972–67982 (2019).

47. Graving, J. M. et al. DeepPoseKit, a software toolkit for fast and robust animal pose estimation using deep learning. Elife 10.7554/eLife.47994 (2019) doi:10.7554/eLife.47994.

48. Roosjen, P. P. J., Kellenberger, B., Kooistra, L., Green, D. R. & Fahrentrapp, J. Deep learning for automated detection of Drosophila suzukii: potential for UAV-based monitoring. Pest Manag Sci 76, 2994–3002 (2020).

49. Mearns, D. S., Donovan, J. C., Fernandes, A. M., Semmelhack, J. L. & Baier, H. Deconstructing Hunting Behavior Reveals a Tightly Coupled Stimulus-Response Loop. Current Biology 30, 54–69.e9 (2020).

50. Layana Castro, P. E., Puchalt, J. C. & Sánchez-Salmerón, A. J. Improving skeleton algorithm for helping Caenorhabditis elegans trackers. Sci Rep 10, (2020).

51. Banerjee, A., Wu, S., Cheng, L. & Aw, S. S. Fully automated leg tracking in freely moving insects using feature learning leg segmentation and tracking (FLLIT). Journal of Visualized Experiments 2020, (2020).

52. Kukieattikool, P. et al. Improvements to the Aedes larvae mobile detection system. Maejo International Journal of Science and Technology 14, 195–208 (2020).

53. Seong, K. H., Matsumura, T., Shimada-Niwa, Y., Niwa, R. & Kang, S. The drosophila individual activity monitoring and detection system (Diamonds). Elife 9, 1–41 (2020).

54. Kasinathan, T., Singaraju, D. & Uyyala, S. Insect classification and detection in field crops using modern machine learning techniques. Information Processing in Agriculture 8, 446–457 (2020).

55. Karashchuk, P. et al. Anipose: A toolkit for robust markerless 3D pose estimation. Cell Rep 36, (2021).

56. Ong, S.-Q., Ahmad, H. & Majid, A. H. A. Development of a deep learning model from breeding substrate images: a novel method for estimating the abundance of house fly (Musca domestica L.) larvae. Pest Manag Sci 77, 5347–5355 (2021).

57. García Garví, A., Puchalt, J. C., Layana Castro, P. E., Navarro Moya, F. & Sánchez-Salmerón, A.-J. Towards Lifespan Automation for Caenorhabditis elegans Based on Deep Learning: Analysing Convolutional and Recurrent Neural Networks for Dead or Live Classification. Sensors 21, (2021).

58. Gosztolai, A. et al. LiftPose3D, a deep learning-based approach for transforming two-dimensional to three-dimensional poses in laboratory animals. Nat Methods 18, 975–981 (2021).

59. Hebert, L., Ahamed, T., Costa, A. C., O’Shaughnessy, L. & Stephens, G. J. WormPose: Image synthesis and convolutional networks for pose estimation in C. elegans. PLoS Comput Biol 17, (2021).

60. Ahmad, M. N., Shariff, A. R. M., Aris, I. & Abdul Halin, I. A Four Stage Image Processing Algorithm for Detecting and Counting of Bagworm, Metisa plana Walker (Lepidoptera: Psychidae). Agriculture 11, (2021).

61. Bjerge, K., Nielsen, J., Sepstrup, M., Helsing-Nielsen, F. & Hoye, T. An automated light trap to monitor moths (Lepidoptera) using computer vision-based tracking and deep learning. Sensors 113, E1082–E1088 (2021).

62. Michels, T., Berh, D. & Jiang, X. An RJMCMC-Based Method for Tracking and Resolving Collisions of Drosophila Larvae. IEEE/ACM Trans. Comput. Biol. Bioinformatics 16, 465–474 (2019).

63. Bodenstein, C., Götz, M., Jansen, A., Scholz, H. & Riedel, M. Automatic Object Detection Using DBSCAN for Counting Intoxicated Flies in the FLORIDA Assay. in 2016 15th IEEE International Conference on Machine Learning and Applications (ICMLA) 746–751 (2016). doi:10.1109/ICMLA.2016.0133.

64. Kesvarakul, R., Chianrabutra, C. & Chianrabutra, S. Baby Shrimp Counting via Automated Image Processing. in Proceedings of the 9th International Conference on Machine Learning and Computing 352–356 (Association for Computing Machinery, 2017). doi:10.1145/3055635.3056652.

65. Almarinez, J. & Hernandez, A. Classifying Mangrove Crub Images for Growth Stages Detection and Monitoring. in 2019 *International Conference on Sensing, Diagnostics*, Prognostics, and Control (SDPC) 710–713 (2019). doi:10.1109/SDPC.2019.00134.

66. Sharma, K., Gold, M., Zurbruegg, C., Leal-Taixé, L. & Wegner, J. D. HistoNet: Predicting size histograms of object instances. in 2020 IEEE Winter Conference on Applications of Computer Vision (WACV) 3626–3634 (2020). doi:10.1109/WACV45572.2020.9093484.

67. Suarez, A. et al. Pest detection and classification to reduce pesticide use in fruit crops based on deep neural networks and image processing. in 2021 XIX WORKSHOP ON INFORMATION PROCESSING AND CONTROL (RPIC) (2021). doi:10.1109/RPIC53795.2021.9648485.

68. Armalivia, S., Zainuddin, Z., Achmad, A. & Wicaksono, M. A. Automatic Counting Shrimp Larvae Based You Only Look Once (YOLO). in 2021 International Conference on Artificial Intelligence and Mechatronics Systems (AIMS) 1–4 (2021). doi:10.1109/AIMS52415.2021.9466058.

69. Saeed, M. S., Nazreen, S. F., Ullah, S. S. S. A., Rinku, Z. F. & Rahman, M. A. Detection of Mosquito Larvae Using Convolutional Neural Network. in 2021 *2nd International Conference on Robotics*, Electrical and Signal Processing Techniques (ICREST) 478– 482 (2021). doi:10.1109/ICREST51555.2021.9331235.

70. Gao, M., Ma, S., Zhao, L. & Zhang, Y. Soybean Pest Recognition Based on YOLOv4 and Graph Cut Algorithm. in 2021 IEEE 3rd International Conference on Frontiers Technology of Information and Computer (ICFTIC) 212–216 (2021). doi:10.1109/ICFTIC54370.2021.9647100.

71. Bentzur, A., Ben-Shaanan, S., Benishou, J. & Shohat-Ophir, G. Social interaction and network structure in groups of Drosophila males are shaped by prior social experience and group composition. BioRxiv 10.1101/2020.03.19.995837.ABSTRACT (2020) doi:10.1101/2020.03.19.995837.ABSTRACT.

72. Li, S. et al. Pose estimation and tracking dataset for multi-animal behavior analysis on the China Space Station. Sci Data 12, (2025).

73. Yang, Q. et al. Reynolds rules in swarm fly behavior based on KAN transformer tracking method. Sci Rep 15, (2025).

74. Arame, M., et al. Detection of Leaf Miner Infestation in Chickpea Plants Using Hyperspectral Imaging in Morocco. Agronomy 15, (2025).

75. Zhang, G. et al. Efficient one-stage detection of shrimp larvae in complex aquaculture scenarios. Artificial Intelligence in Agriculture 15, 338–349 (2025).

76. Dang, T.-H., Dang, N.-H. & Tran, V.-T. Shrimp larvae detection and counting in aquaculture using multiscale feature fusion networks. Comput Electron Agric 233, (2025).

77. Dolata, P., Majewski, P., Lampa, P., Ziȩba, M. & Reiner, J. Amodal Instance Segmentation for Mealworm Growth Monitoring Using Synthetic Training Images. IEEE Access 13, 52157–52175 (2025).

78. Zhan, W., Chen, W. & Pan, Y. Automated Recognition and Analysis of Escape Response in Caenorhabditis elegans. IEEE Transactions on Computational Biology and Bioinformatics https://ieeexplore.ieee.org/abstract/document/10930813/ (2025).

79. McKenzie-Smith, G. C., Wolf, S. W., Ayroles, J. F. & Shaevitz, J. W. Capturing continuous, long timescale behavioral changes in Drosophila melanogaster postural data. PLoS Comput Biol 21, e1012753 (2025).

80. Bates, K., Le, K. & Lu, H. Deep learning for robust and flexible tracking in behavioral studies for C. elegans. PLoS Comput Biol 18, (2022).

81. Pereira, T. D. et al. SLEAP: A deep learning system for multi-animal pose tracking. Nat Methods 4, 978–990 (2022).

82. Ravan, A., Feng, R., Gruebele, M. & Chemla, Y. R. Rapid automated 3-D pose estimation of larval zebrafish using a physical model-trained neural network. PLoS Comput Biol 19, (2023).

83. Gore, S. V, Kakodkar, R., Hernandez, T. D. R., Edmister, S. T. & Creton, R. Zebrafish Larvae Position Tracker (Z-LaP Tracker): a high-throughput deep-learning behavioral approach for the identification of calcineurin pathway-modulating drugs using zebrafish larvae. Sci Rep 13, (2023).

84. Keleş, M. F., et al. FlyVISTA, an Integrated Machine Learning Platform for Deep Phenotyping of Sleep in Drosophila. BioRxiv 10.1101/2023.10.30.564733 (2024) doi:10.1101/2023.10.30.564733.

85. Gendelev, L. et al. Deep phenotypic profiling of neuroactive drugs in larval zebrafish. Nat Commun 15, (2024).

86. Melis, J. M., Siwanowicz, I. & Dickinson, M. H. Machine learning reveals the control mechanics of an insect wing hinge. Nature 628, (2024).

87. Zhang, J. et al. Deep Learning for Microfluidic-Assisted Caenorhabditis elegans Multi-Parameter Identification Using YOLOv7. Micromachines (Basel) 14, (2023).

88. Bian, A., Jiang, X., Berh, D. & Risse, B. Resolving Colliding Larvae by Fitting ASM to Random Walker-Based Pre-Segmentations. IEEE/ACM Trans. Comput. Biol. Bioinformatics 18, 1184–1194 (2019).

89. Al-Marzouqi, A. H. & Arabi, A. A. A Comparative Analysis of the Performance of Leading Countries in Conducting Artificial Intelligence Research. Hum Behav Emerg Technol 2024, 1689353 (2024).

90. The fastest-rising institutions by country for scientific research These 10 countries top the ranks in chemistry research. Preprint at 10.1038/nature-index/news/the-fastest-rising-institutions-by-country-for-scientific-research (2020).

91. Kang, S.-H., Jeong, I.-S. & Lim, H.-S. A deep learning-based biomonitoring system for detecting water pollution using Caenorhabditis elegans swimming behaviors. Ecol Inform 80, (2024).

92. Galimov, E. & Yakimovich, A. A tandem segmentation-classification approach for the localization of morphological predictors of C. elegans lifespan and motility. Aging 14, 1665–1677 (2022).

93. Obasekore, H., Fanni, M., Ahmed, S. M., Parque, V. & Kang, B.-Y. Agricultural Robot-Centered Recognition of Early-Developmental Pest Stage Based on Deep Learning: A Case Study on Fall Armyworm (Spodoptera frugiperda). Sensors 23, (2023).

94. Garcia-Garvi, A., Layana-Castro, P. E., Puchalt, J. C. & Sanchez-Salmeron, A.-J. Automation of Caenorhabditis elegans lifespan assay using a simplified domain synthetic image-based neural network training strategy. Comput Struct Biotechnol J 21, 5049–5065 (2023).

95. Costa, C. S., et al. Counting tilapia larvae using images captured by smartphones. Smart Agricultural Technology 4, (2023).

96. Costa, C., Zanoni, V. & … L. C. Deep learning applied in fish reproduction for counting larvae in images captured by smartphone. Aquac Eng 97, (2022).

97. Manduca, G. et al. Learning algorithms estimate pose and detect motor anomalies in flies exposed to minimal doses of a toxicant. iScience 26, (2023).

98. Zhang, C., Bracke, M., Torres, R. da S. & Gansel, L. C. Rapid detection of salmon louse larvae in seawater based on machine learning. Aquaculture 592, (2024).

99. Jang, G., Yeo, W., Park, M. & Park, Y. G. RT-CLAD: Artificial Intelligence-Based Real-Time Chironomid Larva Detection in Drinking Water Treatment Plants. Sensors 24, (2024).

100. Pham, T. D. Classification of Caenorhabditis Elegans Locomotion Behaviors With Eigenfeature-Enhanced Long Short-Term Memory Networks. IEEE/ACM Trans Comput Biol Bioinform 20, 206–216 (2023).

101. Genaev, M. A. et al. Classification of Fruit Flies by Gender in Images Using Smartphones and the YOLOv4-Tiny Neural Network. Mathematics 10, (2022).

102. Nguyen, M., Roman, G. W. & Soibam, B. Drosophila genotypes can be predicted from their exploration locomotive trajectories using supervised machine learning. Behavioural Processes 212, (2022).

103. Bigge, J., Ogueta, M., Garcia, L. & Risse, B. The Hatching-Box: A Novel System for Automated Monitoring and Quantification of Drosophila melanogaster Developmental Behavior. ArXiv https://arxiv.org/pdf/2411.15390v2 (2024).

104. McClanahan, P. D., Golinelli, L., Le, T. A. & Temmerman, L. Automated scoring of nematode nictation on a textured background. PLoS One 18, (2023).

105. Banerjee, S. C., Khan, K. A. & Sharma, R. Deep-worm-tracker: Deep learning methods for accurate detection and tracking for behavioral studies in C. elegans. Appl Anim Behav Sci 266, (2023).

106. Weng, J., Lin, Q., Chen, S. & Chen, W. Method for Live Determination of Caenorhabditis Elegans Based on Deep Learning and Image Processing. in 2024 *IEEE International Conference on Advanced Information, Mechanical Engineering*, Robotics and Automation (AIMERA) 62–68 (2024). doi:10.1109/AIMERA59657.2024.10735706.

107. Green, A. J. et al. Deep autoencoder-based behavioral pattern recognition outperforms standard statistical methods in high-dimensional zebrafish studies. PLoS Comput Biol 20, (2023).

108. Widrick, J. J. et al. High resolution kinematic approach for quantifying impaired mobility of dystrophic zebrafish larvae. Biorxiv 10.1101/2024.12.05.627004 (2024) doi:10.1101/2024.12.05.627004.

109. Si, G., Zhou, F., Zhang, Z. & Zhang, X. Tracking Multiple Zebrafish Larvae Using YOLOv5 and DeepSORT. in 2022 8th International Conference on Automation, Robotics and Applications, ICARA 2022 228–232 (Institute of Electrical and Electronics Engineers Inc., 2022). doi:10.1109/ICARA55094.2022.9738556.

110. Long, W. et al. An improved shrimp larvae counting algorithm based on HRNET. in 5th International Conference on Artificial Intelligence and Advanced Manufacturing (AIAM 2023) vol. 2023 98–104 (2023).

111. Zhang, G., Shen, Z., Li, D., Zhong, P. & Chen, Y. CAGNet: an improved anchor-free method for shrimp larvae detection in intensive aquaculture. Aquaculture International 32, 6153–6175 (2024).

112. Zhou, C. et al. Counting, locating, and sizing of shrimp larvae based on density map regression. Aquaculture International 32, 3147–3168 (2023).

113. Tiyapunjanit, P., Siammai, T., Klaythong, K., Jantrachotechatchawan, C. & Duangrattanalert, K. VannameiVision: An Optimized Probabilistic Deep Learning for Susceptible Shrimp Larvae Detection. in 2023 3rd International Conference on Emerging Smart Technologies and Applications (eSmarTA) 1–7 (2023). doi:10.1109/eSmarTA59349.2023.10293495.

114. Surya, A., Peral, D. B., VanLoon, A. & Rajesh, A. A Mosquito is Worth 16x16 Larvae: Evaluation of Deep Learning Architectures for Mosquito Larvae Classification. ArXiv https://arxiv.org/pdf/2209.07718 (2022).

115. Abedin, R., Nakib, A. Al, Islam, T. & Islam, M. E. An Improved Transfer Learning Based Larvae Detection and Classification Using Densenet121. in 2023 26th International Conference on Computer and Information Technology (ICCIT) 1–6 (2023). doi:10.1109/ICCIT60459.2023.10441505.

116. Jahangir, R. et al. A Conceptual Framework of an Automated Mosquito Control in Drainage Systems for Combating Dengue in Bangladesh. in 2023 26th International Conference on Computer and Information Technology (ICCIT) 1–6 (2023). doi:10.1109/ICCIT60459.2023.10441613.

117. Javed, N., López-Denman, A. J., Paradkar, P. N. & Bhatti, A. LarvaeCountAI: a robust convolutional neural network-based tool for accurately counting the larvae of Culex annulirostris mosquitoes. Acta Trop 260, 107468 (2024).

118. Lee, S.-H. & Gao, G. A Study on Pine Larva Detection System Using Swin Transformer and Cascade R-CNN Hybrid Model. Applied Sciences 13, (2023).

119. Fawzy, A., Atef, M. & Mohamed, A. Behavioral Analysis of Insects Pollination Effects: A Precision Agriculture Approach. in 2024 *Intelligent Methods*, Systems, and Applications (IMSA) 93–97 (2024). doi:10.1109/IMSA61967.2024.10652623.

120. Majewski, P., Lampa, P., Burduk, R. & Reiner, J. End-to-end Solution for Tenebrio Molitor Rearing Monitoring with Uncertainty Estimation and Domain Shift Detection. in 2024 IEEE/CVF Conference on Computer Vision and Pattern Recognition Workshops (CVPRW) 5498–5507 (2024). doi:10.1109/CVPRW63382.2024.00559.

121. Gunes, H. & Gungormus, A. Identification of honey bee (Apis mellifera) larvae in the hive with faster R-CNN for royal jelly production. J Apic Res 61, 338–345 (2022).

122. Nwoko, I., Ullah, M., Bais, A., Shaun, S. & Wist, T. Monitoring Wheat Midge Populations using CNNs on White Sticky Cards of Pheromone Traps in Field Settings. in 2023 IEEE Canadian Conference on Electrical and Computer Engineering (CCECE) 65–70 (2023). doi:10.1109/CCECE58730.2023.10288734.

123. Liu, P., He, X., Zhao, K., Li, W. & Huang, B. Physiological state recognition model of small silkworm based on improved YOLOv5. Sci Prog 107, (2024).

124. Ahmad, M. N., Shariff, A. R. M. & Moslim, R. The functionality and features of AI automated detector and counter, Oto-BaCTM for bagworm census in oil palm plantation. Indian Journal of Engineering and Material Sciences 31, 914–921 (2024).

125. Deng, S.-C., Lin, H.-J., Chung, P.-L., Liu, A.-C. & Jiang, J.-A. Biological Trajectory Prediction of Beet Armyworm Larva Based on Computer Vision and Time-Series Forecasting Model. in 2024 IEEE International Conference on Evolving and Adaptive Intelligent Systems (EAIS) 1–7 (2024). doi:10.1109/EAIS58494.2024.10570015.

126. Rujito et al. Enhancing Larval Classification Accuracy Through Hyperparameter Optimization in ResNet50 with Three Different Optimizers. in 2023 *International Conference on Modeling & E-Information Research*, Artificial Learning and Digital Applications (ICMERALDA) 56–61 (2023). doi:10.1109/ICMERALDA60125.2023.10458194.

127. Yin, C., Liu, X., Zhang, X., Wang, S. & Su, H. Long 3D-POT: A Long-Term 3D Drosophila-Tracking Method for Position and Orientation with Self-Attention Weighted Particle Filters. Applied Sciences 14, (2024).

128. Zhao, X. et al. A review of convolutional neural networks in computer vision. Artif Intell Rev 57, 1–43 (2024).

129. Mienye, I. D., Swart, T. G. & Obaido, G. Recurrent Neural Networks: A Comprehensive Review of Architectures, Variants, and Applications. Information 2024, *Vol.* 15, *Page 517* 15, 517 (2024).

130. Van Houdt, G., Mosquera, C. & Nápoles, G. A review on the long short-term memory model. Artif Intell Rev 53, 5929–5955 (2020).

131. Berahmand, K., Daneshfar, F., Salehi, E. S., Li, Y. & Xu, Y. Autoencoders and their applications in machine learning: a survey. Artif Intell Rev 57, 1–52 (2024).

132. Nerella, S. et al. Transformers and large language models in healthcare: A review. Artif Intell Med 154, (2024).

133. Kimble, J. & Nüsslein-Volhard, C. The great small organisms of developmental genetics: Caenorhabditis elegans and Drosophila melanogaster. Dev Biol 485, 93– 122 (2022).

134. Pastorino, P., Prearo, M. & Barceló, D. Ethical principles and scientific advancements: In vitro, in silico, and non-vertebrate animal approaches for a green ecotoxicology. Green Analytical Chemistry 8, 100096 (2024).

135. Uddin, S. & Lu, H. Confirming the statistically significant superiority of tree-based machine learning algorithms over their counterparts for tabular data. PLoS One 19, e0301541 (2024).

136. Talaei Khoei, T., Ould Slimane, H. & Kaabouch, N. Deep learning: systematic review, models, challenges, and research directions. Neural Comput Appl 35, 23103–23124 (2023).

137. Alloghani, M., Al-Jumeily, D., Mustafina, J., Hussain, A. & Aljaaf, A. J. A Systematic Review on Supervised and Unsupervised Machine Learning Algorithms for Data Science. 3–21 (2020) doi:10.1007/978-3-030-22475-2_1.

138. Redmon, J., Divvala, S., Girshick, R. & Farhadi, A. You Only Look Once: Unified, Real-Time Object Detection. Proceedings of the IEEE Computer Society Conference on Computer Vision and Pattern Recognition 2016-December, 779–788 (2015).

139. Tomassini, S., Ali Akber Dewan, M., Liaqat Ali, M. & Zhang, Z. The YOLO Framework: A Comprehensive Review of Evolution, Applications, and Benchmarks in Object Detection. Computers 2024, *Vol.* 13, *Page 336* 13, 336 (2024).

140. McKenzie-Smith, G. C., Wolf, S. W., Ayroles, J. F. & Shaevitz, J. W. Capturing continuous, long timescale behavioral changes in Drosophila melanogaster postural data. ArXiv http://www.ncbi.nlm.nih.gov/pubmed/37731659 (2023).

