## Supplementary material for "Systematic Review of Artificial Intelligence use in behavioral analysis of invertebrate and larval model organisms: Methods, Applications and Future Recommendations": File S1

| Model organism | **name:** Galleria mellonella |
| --- | --- |
|  | **developmental stage:** larvae |
|  | **total number of individuals investigated:** 35 |
|  | **genotype:** wild-type |
|  | **Investigated conditions:** cytotoxicity of essential oil x |
| Input data | **type:** video |
|  | **camera model:** Canon 80D digital SLR |
|  | **recording speed (fps):** 24 |
|  | **Recording time:** 5h |
|  | **recording/image resolution (px):** 1048x1048 |
|  | **color format:** monochrome |
|  | **Total number:** 2,000 |
|  | **Lightning conditions:** Red light LED illumination from below |
| Model | **Preprocessing steps:** adaptive thresholding, PCA |
|  | **Method:** DL |
|  | **Task:** classification |
|  | **Model name:** YOLOv8 |
|  | **Input size:** 524x524px |
|  | **Train/valid/test split:** 80:10:10 |
| Model architecture (if applicable) | **Training time (h):** 3 |
|  | **Training epoch:** 150 |
|  | **Learning rate:** 0.001 |
|  | **Optimizer:** Adam |
|  | **Number of layers:** 50 |
|  | **Backbone:** ResNet50 |
| Evaluation metrics: | **Classification:**  Accuracy:  mAP50:  Recall:  F1-score:  Specificity: |
|  | **Detection:**  IoU:  Precision:  Recall:  F1-score: |
|  | **Keypoints detection:**  Percentage of correct keypoints (PCKh@0.5):  Object keypoint similarity: |
|  | **Tracking:**  DetA:  MOTA:  MOTP:  IDF1: |
